# The zebrafish model tackles anti-P2Y_12_ variability in humans: a translational approach

**DOI:** 10.1101/2025.09.21.676872

**Authors:** Paulina Ciepla, Richard J Fish, Séverine Nolli, Sylvie Dunoyer-Geindre, Julia Charlon-Gay, Jean-Luc Reny, Marguerite Neerman-Arbez, Pierre Fontana

## Abstract

Antiplatelet drugs are a pillar in the treatment strategy to prevent ischemic events in cardiovascular patients. However, the action of existing therapies is nonuniform and the causative mechanism for this variability remains poorly understood. The differences between the microRNA profiles of individuals have been suggested to impact platelet reactivity and treatment outcomes. microRNA-150 (miR-150) has been previously associated, in several clinical reports with platelet function variability and cardiovascular events. Therefore, we initiated our analysis by examining the mechanistic role of miR-150 in platelet function. We employed transgenic zebrafish larvae designed to specifically increase miR-150 expression in thrombocytes. Laser-induced caudal vein injury in these animals resulted in a smaller thrombus and decreased thrombocyte accumulation. RNA sequencing of miR-150-overexpressing thrombocytes identified a downregulated transcript encoding *microtubule associated serine/threonine kinase-like* (*mastl*), a key regulator of P2Y_12_ receptor downstream signaling. Overnight treatment of the transgenic fish with clopidogrel, a P2Y_12_ inhibitor, showed decreased thrombus formation in the control, but not in miR-150-overexpressing animals. This phenotype was reversed by overexpressing mastl. Next, we utilized a VASP phosphorylation assay to show that MASTL-deficiency partially protects human platelets from P2Y_12_ inhibition. Finally, using miRNA:mRNA interaction predictors we identified MASTL-targeting miRNAs in humans – miR-17-5p and miR-106a-5p – which were significantly upregulated in a population of clopidogrel-resistant patients. Our studies therefore support a model where an increase in miR-150 in zebrafish and miR-17-5p or miR-106a-5p in humans results in MASTL downregulation, which decreases VASP phosphorylation in platelets and thus makes them resistant to P2Y_12_ inhibition.

**Key points:**

- microRNA-150 downregulates mastl expression in zebrafish larvae, leading to decreased thrombocyte adhesion upon vascular endothelial injury and resistance to P2Y_12_ inhibition.
- In humans, downregulation of MASTL results in a decreased response to P2Y_12_ antagonists.
- MASTL targeting microRNAs are upregulated in cardiovascular patients with poor on-treatment response to clopidogrel.

## Introduction

Antiplatelet drugs, such as P2Y_12_ inhibitors, are at the forefront of the treatment strategy following an ischemic event. However, their action is nonuniform and can be associated with the recurrence of thrombosis or the development of a bleeding episode.^1^ Currently several methods, including the use of platelet function assays and genetic testing, have been proposed to improve the choice of an appropriate type of P2Y_12_ inhibitor for each patient.^2^ However, these approaches remain controversial, due to lack of convincing evidence supporting such approaches.^2,3^ The CYP2C19 polymorphisms explain approximately 10% of the variability in P2Y_12_ inhibitor response,^4^ leaving the remaining 90% unexplained and suggests that other regulatory mechanisms, including microRNAs, play crucial, yet uncharacterized roles in platelet reactivity and drug responses.MicroRNAs (miRNAs) are small, non-coding RNAs (∼20 nucleotides in length) that are highly abundant in mature platelets.^5^ They function by modulating gene expression through the binding to mRNA targets and inhibiting protein translation, either directly or indirectly, by promoting mRNA degradation.^6,7^ Interestingly, specific miRNAs have been previously linked with some cardiovascular related functions, such as haematopoiesis, vasculature homeostasis, inflammation and platelet reactivity.^6-8^ Additionally, the level of circulating, platelet-derived miRNAs has been correlated with the presence or severity of several cardiovascular diseases (CVDs).^9^ However, the mechanistic regulation of platelet function by specific miRNAs remains largely elusive, although pioneering studies have uncovered some mechanistic insights using human cells, mice and zebrafish.^10-14^Animal models have played a critical role in identifying and characterizing many factors and pathways that initiate, drive or resolve thrombotic processes.^15^ They have also aided in the evaluation of the safety and efficacy of antithrombotic treatments.^15^ Among these, the evolutionarily conserved coagulation pathways and a similar arterial and venous clot architecture, combined with the unique optical transparency, have made zebrafish (*Danio rerio*) a powerful model for studying hemostasis and thrombosis.^16,17^ Their original characteristics enable real-time, live-cell imaging of genetically altered thrombocyte adhesion and aggregation immediately after laser-induced endothelial injury.^18^ Additionally, multiple miRNAs as well as their targets are conserved between zebrafish and humans, making it a valuable platform for translational research.^19^ This model was indeed successfully used to study the role of various coagulation and anticoagulant factors,^20-23^ glycoproteins and metalloproteases in hemostasis,^24-26^ and the role of specific mutations found in fibrinogen disorders.^27-29^ Here, we focused our attention on the role of miR-150 in thrombocyte regulation. This small miRNA is abundant in platelets and has been associated with platelet reactivity and cardiovascular events in humans.^30-34^ Additionally, the structure and sequence of miR-150 is evolutionarily conserved among vertebrates, with an identical miRNA seed sequence shared between fish, mice and humans.^35^ Thus, in the present work, we utilized thrombocyte-specific miR-150 overexpression in zebrafish larvae combined with laser-induced endothelial injury, as an entry point to uncover novel regulators of platelet reactivity.

## Methods

Detailed experimental methods can be found in the supplemental materials.

### Generation of transgenic animals

Tol2-mediated transgenesis was utilized to obtain transgenic zebrafish (details are described in the supplemental materials).^36^ If necessary, starting with the F2 generation, adults were mated with *Tg(−6*.*0itga2b:EGFP)* or *Tg(kdrl:EGFP)* fish to perform experiments. Zebrafish line nomenclature: *Tg(−6*.*0itga2b:tagRFP)* from now on will be called *Tg(itga2b:tagRFP*), or the control line; *Tg(−6*.*0itga2b:miR-150-tagRFP)* will be called *Tg(itga2b:miR-150-tagRFP)*.

### Laser-induced vascular endothelial injury and data analysis

Venous thrombosis was assessed in zebrafish larvae with circulating TagRFP fluorescent thrombocytes at 5 days post-fertilization (dpf). The live fish were anesthetized with 170 mg/L tricaine (MS-222, Sigma-Aldrich) and mounted in 0.9% low-melting-point agarose (Sigma-Aldrich) on a glass slide. Laser injuries were performed using Leica DM 6500 microscope (Leica Microsystems) equipped with a HCX PL FLUOTAR L 20x/0.40 corr objective and a crystal laser (wavelength: 355 nm) (more set up details can be found in the supplemental materials). Following the injury, videos of accumulating TagRFP^+^ cells were taken using Leica DFC 360 FX camera and LAS-AF software, at constant exposure (260.25 ms) and with a constant frame rate (263 or 288 ms/frame, experiment dependent) for up to 180 seconds. At approximately 210 seconds, brightfield images of the wound were acquired with a Leica LMD CC7000 camera using LMD software. At least 14 fish per experimental condition were imaged and subsequently analyzed. Details of the data analysis performed are described in the supplemental materials.

### Human platelet generation and the VASP assay

Human hematopoietic stem cells (CD34^+^) were isolated from the buffy coats of healthy adult human donors and cultured for 7 days before inducing their differentiation into megakaryocytes (MKs).^10^ Details can be found in the supplemental materials. At day 15, MKs were taken for mRNA quantitative real-time PCR and western blot analysis, and platelets were taken for the analysis of VASP phosphorylation (all described in the supplemental material).

### Cardiovascular Patients cohort

Stable cardiovascular patients were recruited in the multicenter Antiplatelet Drug Resistance and Ischemic Events (ADRIE) study.^37,38^ The current analysis focused on patients treated with 75 mg of clopidogrel alone or together with 75/100 mg of aspirin. Next, 34 patients with extreme platelet reactivity index (PRI) were matched based on predetermined criteria, known to influence the individual platelet reactivity (**supplemental Table S1**).^37,39,40^ The values for the assessment of low and high platelet reactivity (LPR and HPR) upon clopidogrel treatment were defined using previously described cut-off values as PRI below 16 and above 50, respectively.^2,41,42^

### Quantification of microRNAs in plasma obtained from cardiovascular patients

EDTA-anticoagulated plasma samples from the ADRIE biobank were processed as previously described and are detailed in the supplemental materials.^11,32,37^

## Results

### Strategy for miR-150 overexpression in zebrafish thrombocytes

To study the role of miR-150 in thrombocyte function, we generated a thrombocyte-specific *Danio rerio* miR-150 (dre-miR-150, from now on referred to as miR-150) overexpressing zebrafish transgenic line *Tg(itga2b:miR-150-tagRFP)* and a control line expressing TagRFP only *Tg(itga2b:tagRFP)* (**Figure 1A**). A 30.8±2.7-fold increase in miR-150 expression in thrombocytes (*p* = 0.0075) was detected as compared to the control larvae at 5 days post-fertilization (dpf), with no change of the miR-150 level in the vascular endothelial cells (**Figure 1B**), consistent with the tissue-specific overexpression of this miRNA. Interestingly, the basal level of miR-150 in thrombocytes was lower than that detected in endothelial cells, implying a potential role for this miRNA in endothelia. The overexpression of miR-150 in thrombocytes, however, had no effect on the levels of two other miRNAs that are highly expressed in these cells (dre-miR-126 and dre-miR-223,^11,43^ **supplemental Figure 1A-B**), discriminating against a potential change in the levels of all cellular miRNAs.

**Figure 1.**
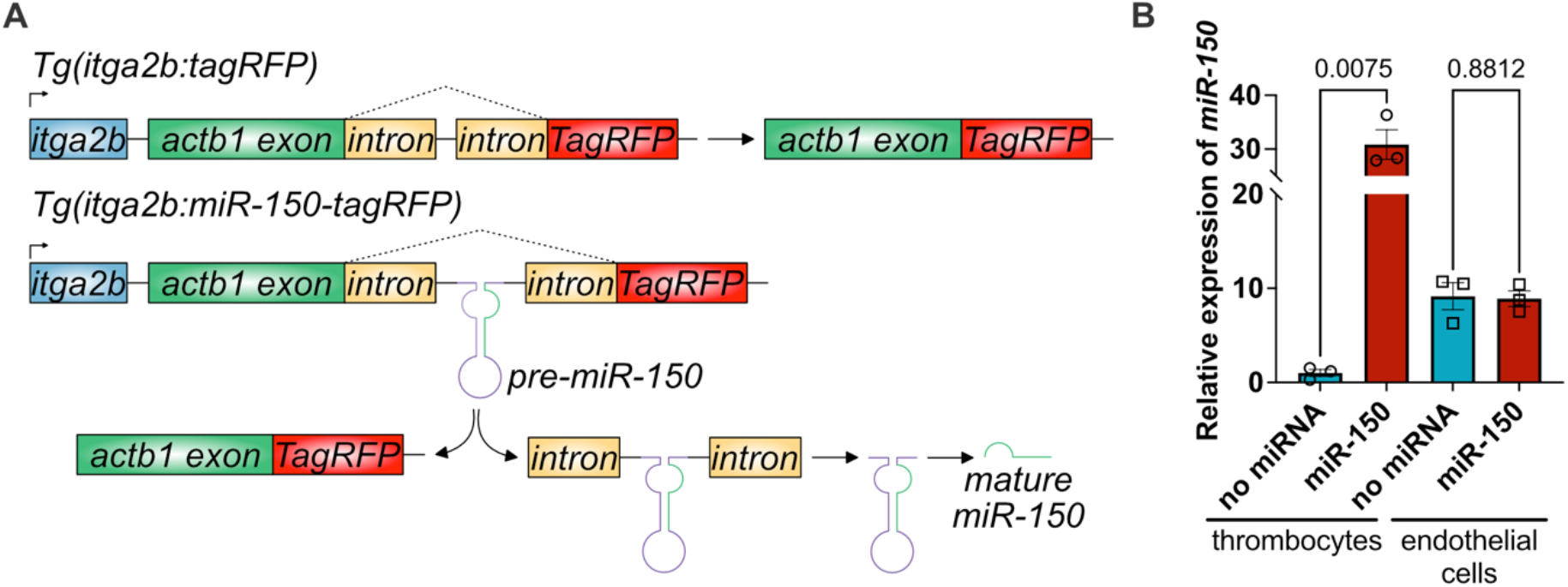
Generation of a thrombocyte-specific, miR-150-overexpressing zebrafish transgenic line. (A) Scheme of the Tol2-itga2b-miR-150-tagRFP construct used to generate the *Tg(itga2b:miR-150-tagRFP)* transgenic zebrafish line and the control line *Tg(itga2b: tagRFP)*. (B) Expression of dre-miR-150 in thrombocytes and vascular endothelial cells of double-transgenic zebrafish larvae (*Tg((itga2b:eGFP),(itga2b:miR-150-tagRFP))* and *Tg((kdrl:EGFP),(itga2b:miR-150-tagRFP))*, respectively, compared to the control lines. The double-positive TagRFP+/GFP+ or GFP+ only cells were collected for the analysis of miRNA levels in different cell types. Results are expressed as fold-change relative to control thrombocytes (n = 3). Error bars represent SEM.

### Characterization of miR-150-overexpressing thrombocytes

No difference in the total number of hematopoietic stem and progenitor cells (HSPCs) or mature thrombocytes was shown in 5 dpf larvae, as assessed by crossing the transgenic animals with the *Tg(itga2b:eGFP)* fish, a well-characterized zebrafish line that marks these two cell populations (**supplemental Figure 2A-C**).^44^ To determine the subpopulation of cells that express the transgenes we studied the expression pattern of the cells labelled with tagRFP (TagRFP^+^). There were no additional TagRFP^+^/GFP^−^ cells detected in both the overexpression and control lines, compared to *Tg(itga2b:eGFP)* larvae (**supplemental Figure 2D)**, suggesting that tagRFP expression is limited to the HSPCs and thrombocytes only. However, the level of tagRFP fluorescence varied significantly between the two lines (**supplemental Figure 2D**), most likely due to a position effect, characterized by a differential genomic integration of the transgenes. The tagRFP protein was found in 99.71±0.29% of thrombocytes in the control line and in 89.86±1.49% of thrombocytes in the miR-150-overexpressing line (**supplemental Figure 2E**). Interestingly, the tagRFP^+^ cells were also found in a small population of HSPCs (14.25±2.10% and 8.90±0.80% in the control and the miR-150-overexpressing fish, respectively, **supplemental Figure 2F**), although these cells were not characterized further. To determine the expression pattern of the tagRFP^+^/GFP^+^ cells we imaged the miR-150-overexpressing larvae at 3, 4 and 5 dpf in three regions: aorta/gonad/mesonephros (AGM) and pronephric ducts, caudal hematopoietic tissue (CHT), and thymic lobes, all of which are well-defined areas of zebrafish larvae hematopoiesis (**supplemental Figure 2G-I**, respectively).^45^ The expression of stationary red labeled cells was largely limited to the CHT region, where the larval HSPCs expansion and differentiation occurs.^46^ There were no tagRFP^+^ cells present in thymic lobes at all three developmental timepoints, nor in the AGM and pronephric ducts at the two later stages. A small amount of tagRFP^+^ cells were found in the AGM at 3 dpf in some animals (**supplemental Figure 2G**), which could indicate the possible origin of the first thrombocytes present in the circulation. Finally, no change in the relative size of thrombocytes was observed between the *Tg(itga2b:miR-150-tagRFP)* and *Tg(itga2b:tagRFP)* animals at 5 dpf, as assessed by flow cytometry (**supplemental Figure 2J**). Altogether, our data suggest that the overexpression of miR-150 in zebrafish thrombocytes is specific, does not lead to any global defects in HSPC survival or their differentiation, and does not affect the number or size of thrombocytes in these animals; thus, miR-150 overexpression does not indirectly interfere with thrombus formation.

### miR-150 effects thrombus formation in zebrafish larvae through the downregulation of mastl

The evaluation of the formation of thrombocyte-rich thrombi, within 160 seconds post laser injury, showed a reduction in thrombocyte adhesion in the miR-150-overexpressing zebrafish larvae versus the transgenic controls (36.2±13.6%; *p* = 0.0133; **Figure 2A-B**). Additionally, measurement of the thrombus size approximately 210 seconds post-injury confirmed this observation, with smaller thrombi in the miRNA overexpressing fish being observed (−33.0±15.3%; *p* = 0.0400; **supplemental Figure 3A-B**).Next, RNA sequencing (RNA-seq) analysis was performed on the double fluorescent (TagRFP^+^/GFP^high^) thrombocytes acquired from 5 dpf *Tg((itga2b:miR-150-tagRFP); (itga2b:eGFP))* or from *Tg((itga2b:tagRFP); (itga2b:eGFP))* larvae. Using the R/Bioconductor package edgeR,^47^ we identified 58 differentially expressed genes: 55 upregulated and three downregulated (DEGs, adjusted P > 0.05, **Figure 2C, supplemental Table S2**) in the miRNA overexpressing cells versus the controls. We focused our attention on the downregulated genes, since these could potentially be directly targeted by miR-150. Studying the consensus of the two zebrafish-specific miRNA:mRNA interaction prediction tools: TargetScanFish (release 6.2, https://www.targetscan.org/fish_62/, accession date: 18^th^ March 2024)^48,49^ and DIANA microT-CDS (version 5.0, https://dianalab.e-ce.uth.gr/html/dianauniverse/index.php?r=microT_CDS&threshold=0.3&page=1, accession date: 20^th^ March 2024),^50,51^ we established that only *microtubule associated serine/threonine kinase like* (*mastl*) contains a sequence that matches a dre-miR-150 seed region (7-mer). In accordance with this, our miR-150-overexpressing thrombocytes showed no detectable raw reads mapped to the *mastl* transcript (**supplemental Figure 3C**).

**Figure 2.**
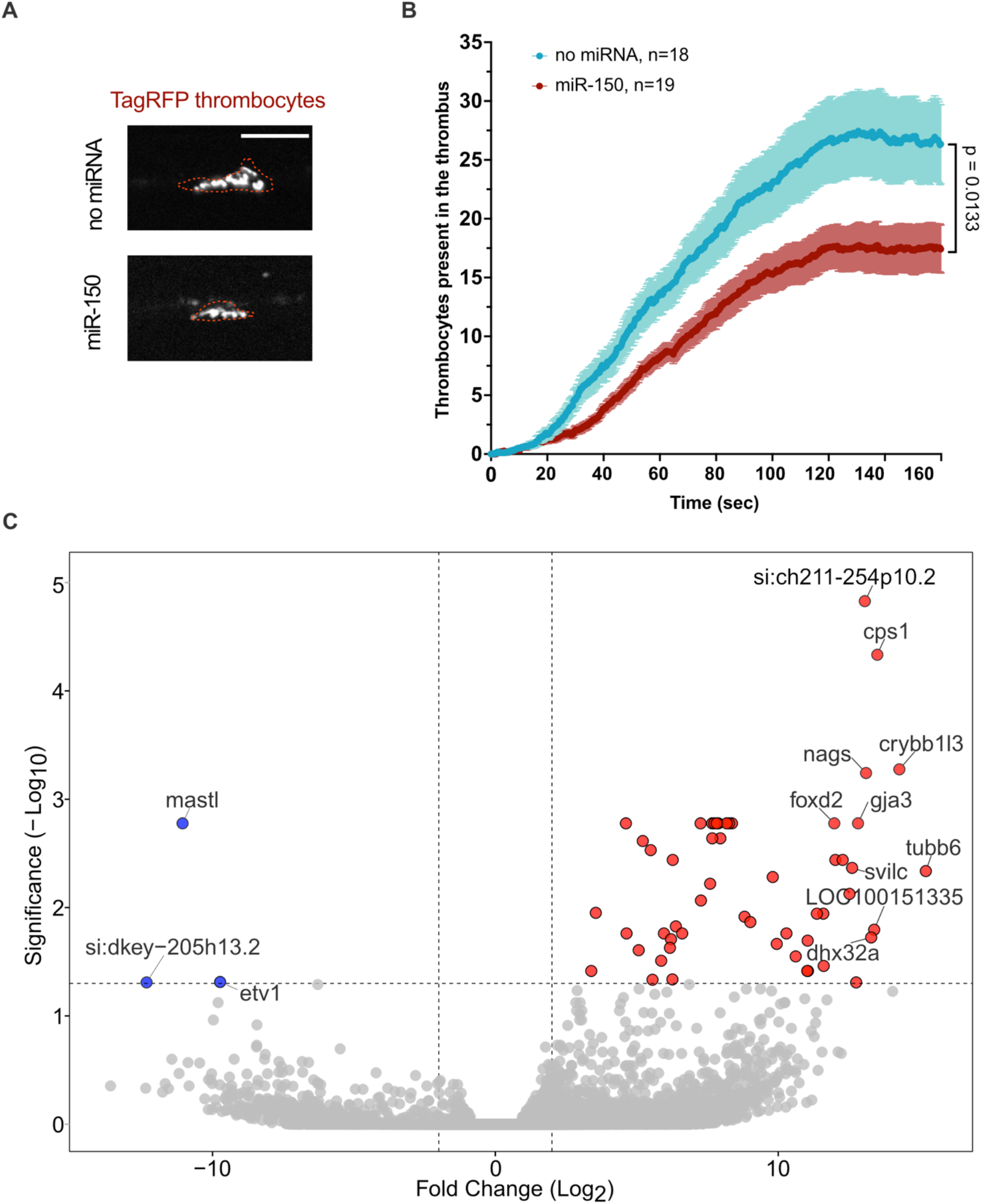
Thrombus formation after a laser-induced vascular endothelial injury in control and miR-150-overexpressing fish and possible miRNA target determination. (A) Representative tagRFP fluorescent image of a thrombus at 150 seconds after laser-induced caudal vein injury in the control and *Tg((itga2b:miR-150-tagRFP))* zebrafish larvae at 5 dpf. Scale bar: 100 μm. (B) Quantification of thrombocyte attachment at the site of laser injury over time in the control: *Tg(itga2b:tagRFP)* (blue; n = 18) and *Tg((itga2b:miR-150-tagRFP))* (red; n = 19) fish at 5 dpf. Error bars represent SEM. (C) Summary of the RNA sequencing data. Volcano plot of negative log_10_ significance *p*-values (with an FDR 5% correction) against log_2_ fold change values, showing differentially expressed genes (DEGs) between miR-150-overexpressing thrombocytes and control. Red dots represent the DEGs with log_2_FC > 2 and significance value <0.05. Blue dots represent the DEGs with log_2_FC < −2 and significance value <0.05. Genes with −2 <log2FC <2 and/or adj. *p*-value >0.05 are shown in grey. Horizontal dashed line marks gene expression with significance value of 0.05; vertical dashed lines represent log_2_FC of 2 (right line) and log_2_FC of −2 (left line). The names of some of the top DEGs are displayed.

### MASTL regulates the response to P2Y_12_ inhibitors but not aspirin in zebrafish

The role of MASTL kinase in platelets is complex and affects many biological pathways, including the ADP-mediated platelet activation, a target of P2Y_12_ inhibitors. Therefore, we hypothesized that the modulation of MASTL by miR-150-overexpression can affect the potential action of P2Y_12_ inhibitors in zebrafish. To test this hypothesis, we performed an overnight treatment of miR-150-overexpressing and control larvae at 4 dpf with clopidogrel (10 µg/mL) or aspirin (180 µg/mL). The successful, on-target use of both clopidogrel and aspirin in zebrafish has previously been described.^11,52-55^ At 5 dpf the larvae were subjected to laser-induced endothelial injury and thrombocyte attachment as well as thrombus size were assessed. The aspirin treatment resulted in a reduction of 44.5±15.7% (*p* = 0.0077) and 66.9±18.1% (*p* = 0.0011) in thrombocyte adhesion post-injury in the control and miR-150-overexpressing fish (**Figure 3A-B, supplemental Figure 4A**). Similarly, the thrombi size decreased by 75.6±23.6% (*p* = 0.0046) and 48.4±17.3% (*p* = 0.0092) in these lines (**supplemental Figure 4B**). Interestingly, clopidogrel treatment resulted in a 41.3±17.3% (*p* = 0.0223) reduction in thrombocyte accumulation post-injury in control animals but had no effect on miR-150-overexpressing fish (**Figure 3A-B, supplemental Figure 4A**). Similarly, the thrombi size after the injury decreased by 52.3±25.3% (*p* = 0.0493) in the control line but was unchanged in the miR-150-overexpressing larvae (**supplemental Figure 4B**). Altogether, these data suggest the successful inhibition of zebrafish thrombocytes with aspirin in both transgenic lines, but the lack of response to clopidogrel only in fish with upregulated miR-150 in their thrombocytes. Next, to prove that this phenotype was at least partially driven by *mastl*, we upregulated its expression in miR-150-overexpressing fish by injection of their zygotes with constructs containing a wild-type (WT) *mastl* coding sequence or a hyperactive *mastl* (K62M) mutant under control of the *itga2b* promoter (**Figure 3C**). The location of the mutation was based on previous literature data and an alignment of *Drosophila*, human and zebrafish protein sequences (**supplemental Figure 5A**).^56,57^ As a negative control we also synthesized a construct where the *mastl* sequence was replaced by blue fluorescent protein (BFP). Next, the injected animals were treated with clopidogrel and subjected to laser injury the following day. As expected, the injection of the negative control construct did not result in a significant difference in thrombocyte accumulation or thrombi size post-injury in the DMSO control and the inhibitor-treated animals (**supplemental Figure 5B-D**). However, *mastl* overexpression resulted in a significant reduction in both thrombocyte accumulation and thrombi size upon treatment by 57.7±18.9% (*p* = 0.0045) and 59.2±14.6% (*p* = 0.0004), respectively (**Figure 3D, supplemental Figure 5C-D**). An even more striking phenotype was observed in fish injected with the *mastl K62M* mutant, where thrombocyte adhesion was reduced by 76.8±13.5% (*p* = <0.0001) and thrombi size by 64.8±14.0 % (*p* = <0.0001; **supplemental Figure 5B-D**). These data suggest that the recovery of mastl expression (both WT and the hyperactive variant) in miR-150-overexpressing thrombocytes can rescue their resistance to P2Y_12_ inhibition.

**Figure 3.**
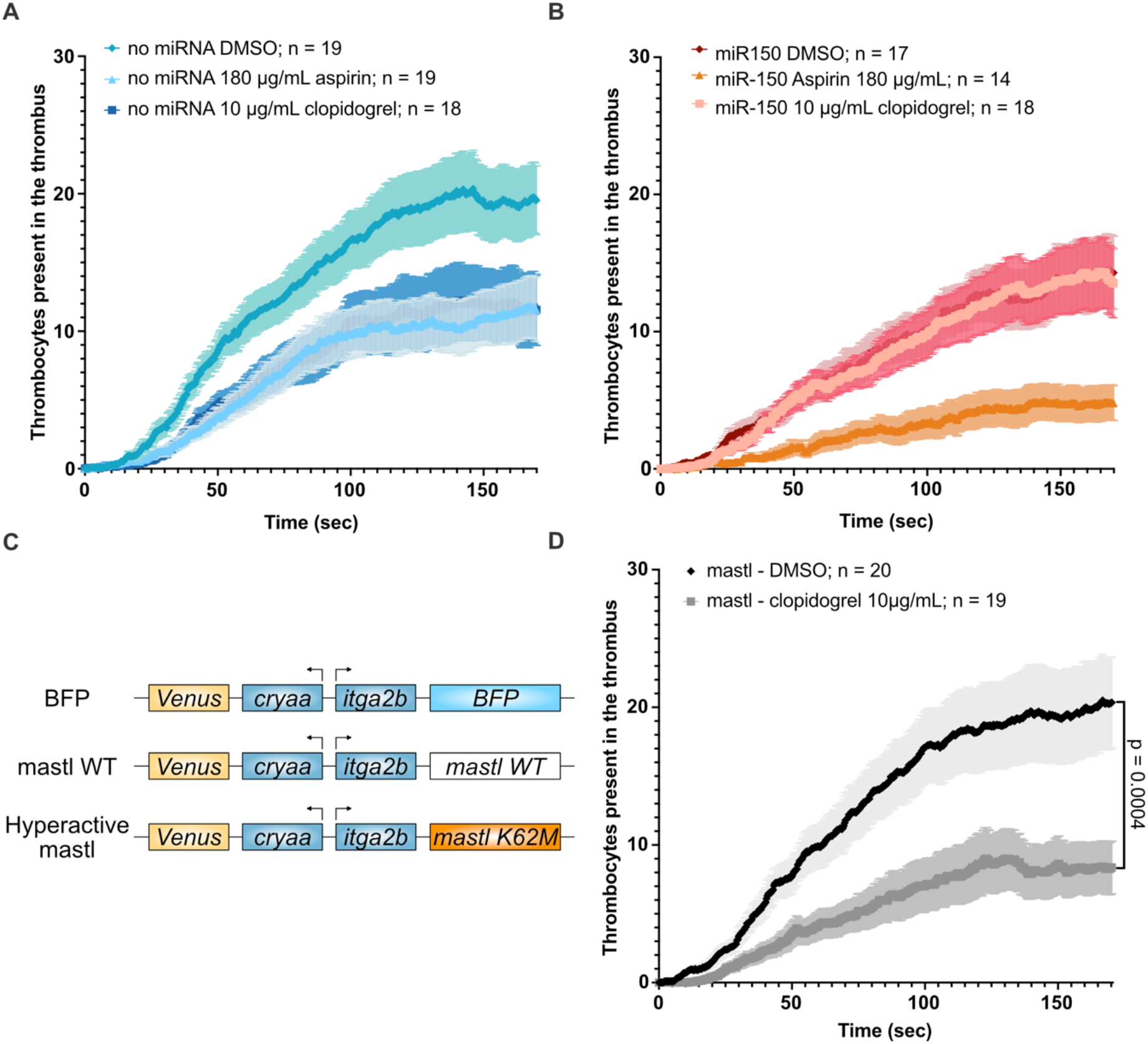
Effect of the selected antiplatelet drug treatment on thrombus characteristics after laser injury in an miR-150-overexpressing line and controls. (A) Quantification of thrombocyte attachment at the site of laser injury over time in *Tg(itga2b:tagRFP)* fish, at 5 dpf, treated overnight with DMSO (n =19), 180 μg/mL aspirin (n =19) or 10 μg/mL clopidogrel (n = 18). (B) Quantification of thrombocyte attachment at the site of laser injury over time in *Tg(itga2b:miR-150-tagRFP)* fish, at 5dpf, treated overnight with DMSO (n =17), 180 μg/mL aspirin (n =14) or 10 μg/mL clopidogrel (n = 18). (C) Scheme of the Tol2-cryaa-Venus-itga2b-mastl (WT and K62M) constructs or of the control construct, Tol2-cryaa-Venus-itga2b-BFP, used to enable a mosaic expression of zebrafish mastl WT, mastl mutant K62M) or BFP specifically in thrombocytes. (D) Quantification of thrombocyte attachment at the site of laser injury over time in *Tg(itga2b:miR-150-tagRFP)* fish, at 5 dpf, injected with the Tol2-cryaa-Venus-itga2b-mastl construct and treated overnight with DMSO (n = 20) or 10 μg/mL clopidogrel (n = 19). Error bars represent SEM.

### Zebrafish thrombocyte reactivity is influenced by mastl expression

This experiment was also used to check the ability of mastl to rescue the reduced thrombus formation phenotype in miR-150-overexpressing fish treated with DMSO only. The injection of the *mastl* construct resulted in an increase in thrombocyte accumulation post-injury by 84.2±33.4 % (*p* = 0.0172) as compared to BFP-injected animals (**supplemental Figure 5C**). The thrombi size after injury showed a tendency to increase (30.0±21.8 %; *p* = 0.1774), although this value did not differ significantly from the negative control (**supplemental Figure 5D**). Interestingly, the *mastl K62M* mutant showed a large and significant increase in the thrombus size after the vein injury (55.2±22.4 %; *p* = 0.0180) but exhibited only a nonsignificant tendency to increase thrombocyte accumulation (32.5±23.0 %; *p* = 0.1655) (**supplemental Figure 5C-D**). This is likely due to a slight increase in the instances of vessel occlusion after the injury in the *mastl K62M* mutant fish versus BFP controls (59.1% and 45.5%, respectively), where upon vein blockage and cessation of blood flow no further thrombocytes can bind to the injury site. Our data indicate that mastl overexpression in miR-150-rich thrombocytes can (at least partially) increase the thrombus size and the number of thrombocytes present within after the laser injury, resembling the control phenotype where the level of miR-150 is lower (*Tg((itga2b:tagRFP)*).

### MASTL regulates the response to P2Y_12_ inhibitors in human cells

Next, we set out to determine whether the decrease of MASTL in platelets derived from human hematopoietic stem cells (CD34^+^)^10,58^ could protect these cells from the effect of P2Y_12_ inhibition. To this end, we nucleofected the megakaryocytes with a small interfering RNA targeting MASTL (siMASTL)^59^ or a scrambled control and these cells were collected after 48 h to assess the decrease of *MASTL* mRNA (72.3±3.3 %; *p* = <0.0001) and protein (59.2±5.8 %; *p* = 0.0002) (**supplemental Figure 6A-C**). Next, we performed a VASP phosphorylation assay using platelets subjected to a preincubation step with cangrelor, a potent P2Y_12_ inhibitor (50 nM), or a water control.^41,42^ The use of an alternative P2Y_12_ inhibitor was necessary due to the requirement of metabolic clopidogrel activation,^60^ unattainable in our *in vitro* cellular system. The platelet reactivity index (PRI) for each sample was calculated to reflect the level of VASP dephosphorylation upon platelet activation with ADP.^41,42^ There were no observed differences in VASP phosphorylation in platelets with reduced levels of MASTL upon treatment with water (**Figure 4A**). The cangrelor preincubation step resulted in a significant reduction in PRI in both cells (64.3±6.3 %; *p* = 0.0002 and 52.3±10.0 %; *p* = 0.0033 in control and siMASTL, respectively), but the index was significantly higher (35.4±10.7 %; *p* = 0.0208) in cells where MASTL was downregulated. This suggests that the decrease in cellular level of MASTL induces (at least partially) platelet resistance to the effects of P2Y_12_ inhibition.

**Figure 4.**
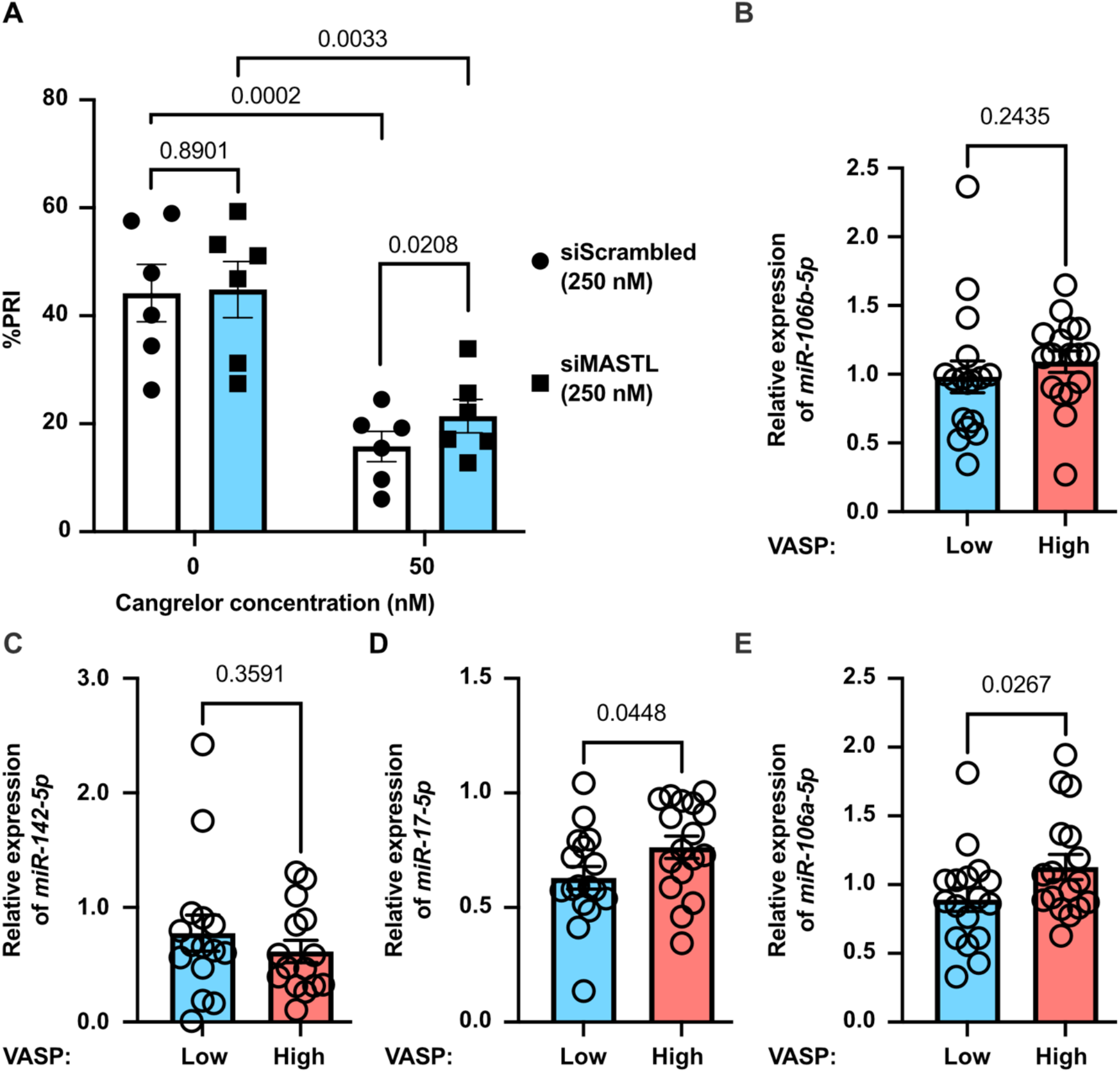
Testing the effect of MASTL deficiency on clopidogrel responses in human platelets and CVD patients. (A) Quantification of the platelet reactivity index (%PRI) in cells derived from megakaryocytes after their nucleofection with scrambled siRNA (control) or *MASTL*-targeting siRNA (siMASTL) and with pretreatment using DMSO or cangrelor (50 nM) before the assay (n = 6). (B-E) Expression of hsa-miR-106-5p (B), hsa-miR-142-5p (C), hsa-miR-17-5p (D) and hsa-miR-106a-5p (E) in plasma obtained from cardiovascular patients treated with clopidogrel (75 mg) with low and high PRI (n = 17, except hsa-miR-142-5p (C): n = 15). Results in B-E are normalized to the stable endogenous miRNAs: hsa-miR-16-5p, hsa-miR-93-5p and hsa-miR-484. Error bars represent SEM.

### Overexpression of MASTL targeting miRNAs in clopidogrel resistant patients

Finally, we tried to determine if the increased expression of miRNAs can influence clopidogrel responsiveness in cardiovascular patients. Using a consensus found between four different online miRNA:mRNA interaction prediction tools (TargetScan; release 8.0; https://www.targetscan.org/vert_80/,^61,62^ PicTar; http://pictar.mdc-berlin.de;^63^ DIANA-microT 2023; https://dianalab.e-ce.uth.gr/microt_webserver/#/,^64^ and miRDB; https://mirdb.org;^65^ (accession date: 25^th^ March 2024)) we discovered that hsa-miR-150-5p is not widely predicted to target the human *MASTL* transcript. However, utilizing these tools, we identified other likely candidate miRNAs: hsa-miR-142-5p and a family of miRNAs that share the seed sequence: AAAGUGC (hsa-miR-17-5p/hsa-miR-106a-5p/hsa-miR-106b-5p) that are expressed in platelets^5,8,13,66,67^ and have previously been correlated with a clopidogrel resistance phenotype in CVD patients.^68,69^ To determine whether the expression of these four miRNAs indeed differs in response to clopidogrel treatment we performed miRNA quantification in plasma samples obtained from 34 stable, symptomatic CVD patients treated with clopidogrel with extreme PRI matched for sex and other factors known to influence clopidogrel response (**supplemental Table S1**).^37,39,40^ After the normalization of the miRNA levels in all 34 samples we determined that the expression of hsa-miR-106b-5p and hsa-miR-142-5p did not differ significantly when comparing the two extreme patient groups (**Figure 4B-C**), however the quantification of hsa-miR-17-5p and hsa-miR-106a-5p showed their significant upregulation in patients with high on-treatment platelet reactivity (121.2±7.8 %; *p* = 0.0448 and 126.7±10.3 %; *p* = 0.0267, respectively) (**Figure 4D-E**), suggesting their role in the poor response to P2Y_12_ inhibition.

## Discussion

Zebrafish has previously emerged as an important *in vivo* model that can uncover unknown genetic regulators of platelet function and provide useful information regarding platelet responses to antiplatelet drugs, all highly relevant to human health.^11,16,20-29,70,71^ In the current study, we used this model to study the influence of transgenic overexpression of miR-150 in zebrafish thrombocytes and determined that its modulation significantly impacts their function. Critically, we first verified that the transgene expression was mostly limited to thrombocytes and not HSPCs, since the *itga2b* promoter (used in our study) can potentially become active in HSPCs (from 27 hours post fertilization),^44,45^ and the miR-150 has previously been shown to influence the expression of *c-myb* and thus alter thrombocyte-erythrocyte linage determination.^35,72^ Next, we determined that this overexpression has no adverse effect on the number of thrombocytes present, their differentiation, their size, or the expression of other miRNAs, suggesting that our strategy was appropriate for studies of platelet function. The utilization of the laser blood-vessel injury assays revealed that miR-150 overexpression results in defective thrombocyte aggregation at the site of endothelial damage and subsequent RNA sequencing pointed to the downregulation of the *microtubule-associated serine/threonine kinase-like* (*mastl*) transcript in these thrombocytes as a potential mechanism for this observed phenotype. *Mastl* gene has previously been shown to regulate platelet number and function.^73-75^ In mice and zebrafish, downregulation of Mastl has been shown to lead to thrombocytopenia.^73,75^ Additionally, the substitution of cytosine to guanidine at nucleotide position 565 in the human *MASTL* gene has also been linked to nonsyndromic autosomal dominant thrombocytopenia (thrombocytopenia-2 (THC2)).^74,75^ In our zebrafish line (*Tg(itga2b:miR-150-tagRFP)*), we avoided the thrombocytopenia phenotype since the modulation of the *mastl* level occurs later in the thrombocyte differentiation process by the experimental design, which ensured almost exclusive thrombocyte-limited transgene expression (TagRFP^+^ cells). In addition to platelet number the lack of Mastl in mice has also been shown to lead to defects in platelet aggregation.^73^ This finding phenocopies our finding in zebrafish where both thrombocyte accumulation and thrombus size are decreased.

Mechanistically, cellular MASTL kinase phosphorylates two small inhibitory proteins: alpha-endosulfine (ENSA) and cAMP-regulated phosphoprotein 19 (ARPP19), which in turn can bind with high affinity to serine/threonine phosphatase (PP2A-B55).^76,77^ This complex leads to the inhibition of PP2A by maintaining it the phosphorylated state (**Figure 5**).^76-78^ In resting platelets active PP2A associates with _αII β III_ integrin, preventing its adhesion to fibrinogen.^79^ In more detail, PP2A is known to counteract the action of major kinases, such as PKC, CMK2 and CK2, and thus influence the activation state of many critical signaling molecules such as: PKCζ, VASP, AKT, ERK, P38 or Syk.^78-83^ This most often leads to their inactivation and to the negative regulation of platelet activation and adhesion. Upon integrin engagement, however, PP2A activity is decreased and integrin-mediated outside-in signaling is unleashed.^79^ Since the downregulation of *mastl*, found in miR-150-overexpressing thrombocytes, should retain PP2A in an active state, it is unsurprising that both zebrafish and mice deficient in Mastl exhibit reduced thrombocyte aggregation and smaller thrombus phenotypes.^73^

**Figure 5.**
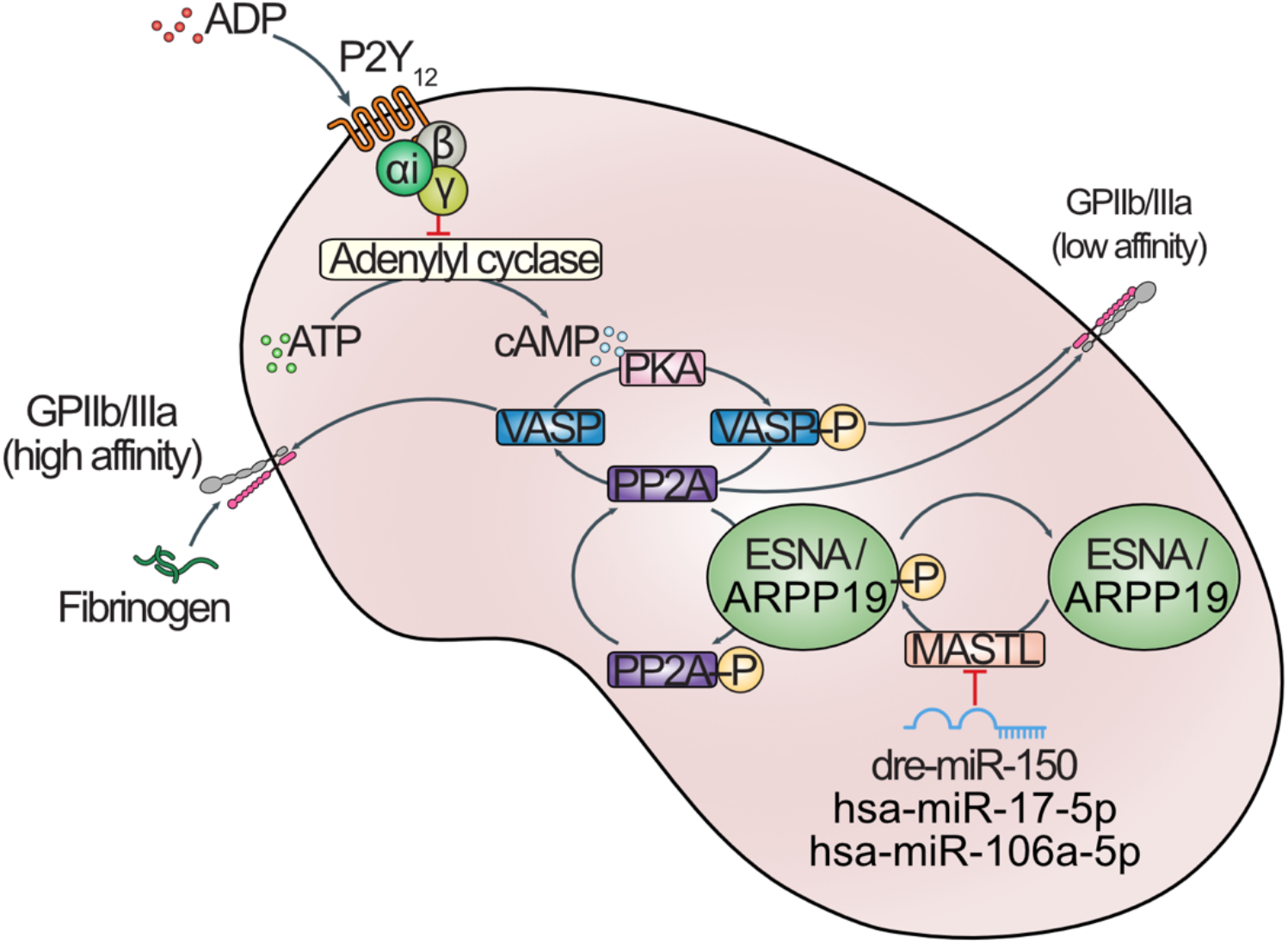
Model representing the role of dre-miR-150, hsa-miR-17-5p and hsa-miR-106a-5p in the regulation of MASTL and platelet reactivity. In platelets the level of MASTL kinase is negatively correlated with the expression of miRNAs (in zebrafish: miR-150 and in human: miR-17-5p and miR-106a-5p). MASTL is responsible for the phosphorylation and thus activation of two kinases: ESNA and ARPP19, which in turn can phosphorylate and inhibit the action of PP2A phosphatase. In the resting platelet, both the direct binding of active PP2A (dephosphorylated form) to GPIIb/IIIa integrin and phosphorylated VASP (VASP-P), keep the GPIIb/IIIa integrin in the closed, inactive conformation, which is characterized by low affinity toward fibrinogen. During platelet activation the direct binding of PP2A to integrin is inhibited. Concurrently, ADP activates the P2Y_12_ receptor, inhibits the action of adenylyl cyclase, thereby reducing the level of cAMP in platelets. This in turn reduces the PKA-mediated phosphorylation of VASP (VASP-P), simultaneously the active PP2A phosphatase dephosphorylates VASP. VASP then activates GPIIb/IIIa integrin and increases its affinity for fibrinogen.

Many of the PP2A targets, including VASP, are components of the ADP-mediated signaling pathway, therefore it could be anticipated that this pathway is adversely affected by the reduction of MASTL levels. Here, MASTL acts as an indirect inhibitor of PP2A,^57,84^ which in turn can dephosphorylate VASP,^79,81^ leading to the activation of_αIIbβIII_ integrin (**Figure 5**).^85^ This pathway is also one of the main targets of many antiplatelet drugs such as: clopidogrel, prasugrel, ticagrelor and cangrelor.^86,87^ These medications target the P2Y_12_ receptor, leading to an increase in the cellular cyclic AMP level and maintaining VASP in an inactive, phosphorylated form, which in turn limits the activation of _αIIbβIII_ integrin and reduces platelet aggregation.^88^ Altogether, we hypothesized that a reduction in MASTL levels can counteract VASP phosphorylation that occurs when the P2Y_12_ antagonist is present. In accordance with this mechanism, our finding has shown that a reduced level of *mastl* decreases the thrombocyte response to clopidogrel, which was successfully rescued by overexpression of *mastl* in these cells. To prove that MASTL also affects the ADP-mediated signaling pathway in human platelets, we reduced MASTL in human platelets, derived from hematopoietic stem cells (CD34^+^), and measured its VASP phosphorylation in the presence or absence of the P2Y_12_ inhibitor cangrelor. In this assay we identified a similar decrease in platelet response to the inhibitor, although this phenotype was less striking than that observed in the zebrafish larvae, most likely due to: i) a large portion of MASTL protein (∼40%) still present in human megakaryocytes and platelets; ii) the markedly different nature of the assays performed in the two model systems; and iii) the use of different inhibitors. To our knowledge this is the first time that MASTL has been directly and mechanistically linked with the ADP-mediated platelet activation pathway. Interestingly, our discovery also provides an additional explanation for interindividual anti-P2Y_12_ drug response variability in humans. Although we could not directly measure the level of MASTL in the platelets of patients of the ADRIE study with high/low on clopidogrel treatment platelet reactivity (due to a lack of appropriate samples), we found that the miRNAs predicted to target MASTL in human cells (miR-17-5p and miR-106a-5p) were differentially expressed in the patients’ plasma when comparing the two groups. These two miRNAs have also been highlighted in independent clinical studies of clopidogrel resistance.^68,69^ A larger study with an extended patient cohort is now required to determine whether these miRNAs could act as biomarkers of the patient’s responses to P2Y_12_ inhibitors.

Taken together our studies emphasize a remarkable conservation in the platelet biology shared between zebrafish and humans, highlighting their use in thrombotic research. However, there are also differences that cannot be overlooked, with the largest limitation in our studies being the lack of the mechanistic details related to the role of miR-150 in platelet function. This miRNA does not target MASTL in human cells, thus, the changes in platelet reactivity observed in the miR-150 related clinical studies^30-32,34^ cannot be directly explained by the modulations of the MASTL level or PP2A activity. Our research also emphasized the need for a new angle of analysis to better understand the reasons behind interindividual anti-P2Y_12_ variability. In contrast to previous whole exome sequencing studies (WES), which failed to identify major determinants of clopidogrel responsiveness,^89,90^ we believe that it is time to invest effort and resources in performing individual miRNA profiling analyses. These small RNAs still play a largely understudied role in platelet biology and cardiovascular diseases, which hinders their potential use as biomarkers of platelet reactivity and treatment outcomes.

## Acknowledgments

The authors would like to thank the members of the Flow Cytometry and Bioimaging core facilities at the University of Geneva for their technical assistance. We would also like to thank the iGE3 Genomics Platform at the University of Geneva (https://ige3.genomics.unige.ch) for performing the RNA sequencing analyses and human plasma miRNA quantification. We also thank blood donors and the production and quality-control staff at Geneva’s blood transfusion center. This study was supported by the Swiss National Science Foundation for Scientific Research (grant # 310030_196864 to P. Fontana and M. Neerman-Arbez) and the Edmond J. SAFRA Fund (unrestricted research grant) to P. Fontana and J.-L. Reny.

## Authorship Contributions

P.C., R.J.F., S.D.G, J.L.R., M.N.A and P.F. designed the study. P.C., S.N. and J.C.G. performed the experiments and analyzed the data. P.C. and P.F. wrote the manuscript’s first draft, and all authors revised the intellectual content and approved the final version.

