## Supplemental Materials for "The zebrafish model tackles anti-P2Y_12_ variability in humans: a translational approach"

### Figure legends

**Sup. Figure 1**

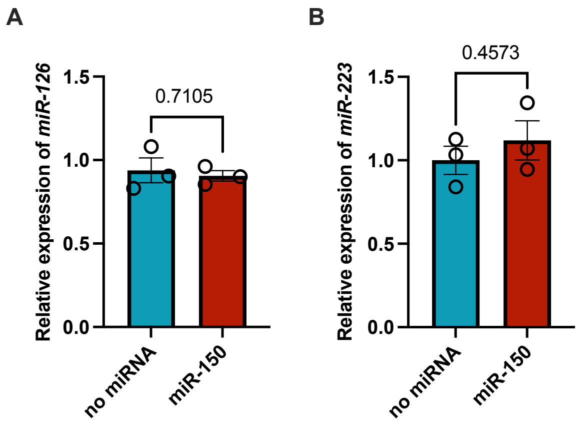


**Supplemental Figure 1. Expression of miRNAs in the thrombocytes of miR-150-overexpressing fish and control fish.** (A) Expression of dre-miR-126 in the thrombocytes of double-transgenic zebrafish line *Tg((itga2b:eGFP),(itga2b:miR-150-tagRFP))* compared to the control line. (B) Expression of dre-miR-223 in the thrombocytes of double-transgenic zebrafish line (*Tg((itga2b:eGFP),(itga2b:miR-150-tagRFP))*) compared to the control line. The double-positive TagRFP/GFP$^+$ cells were collected for the analysis of miRNAs levels. Results are expressed as fold-change relative to control thrombocytes (n = 3). Error bars represent SEM.

Sup. Figure 2

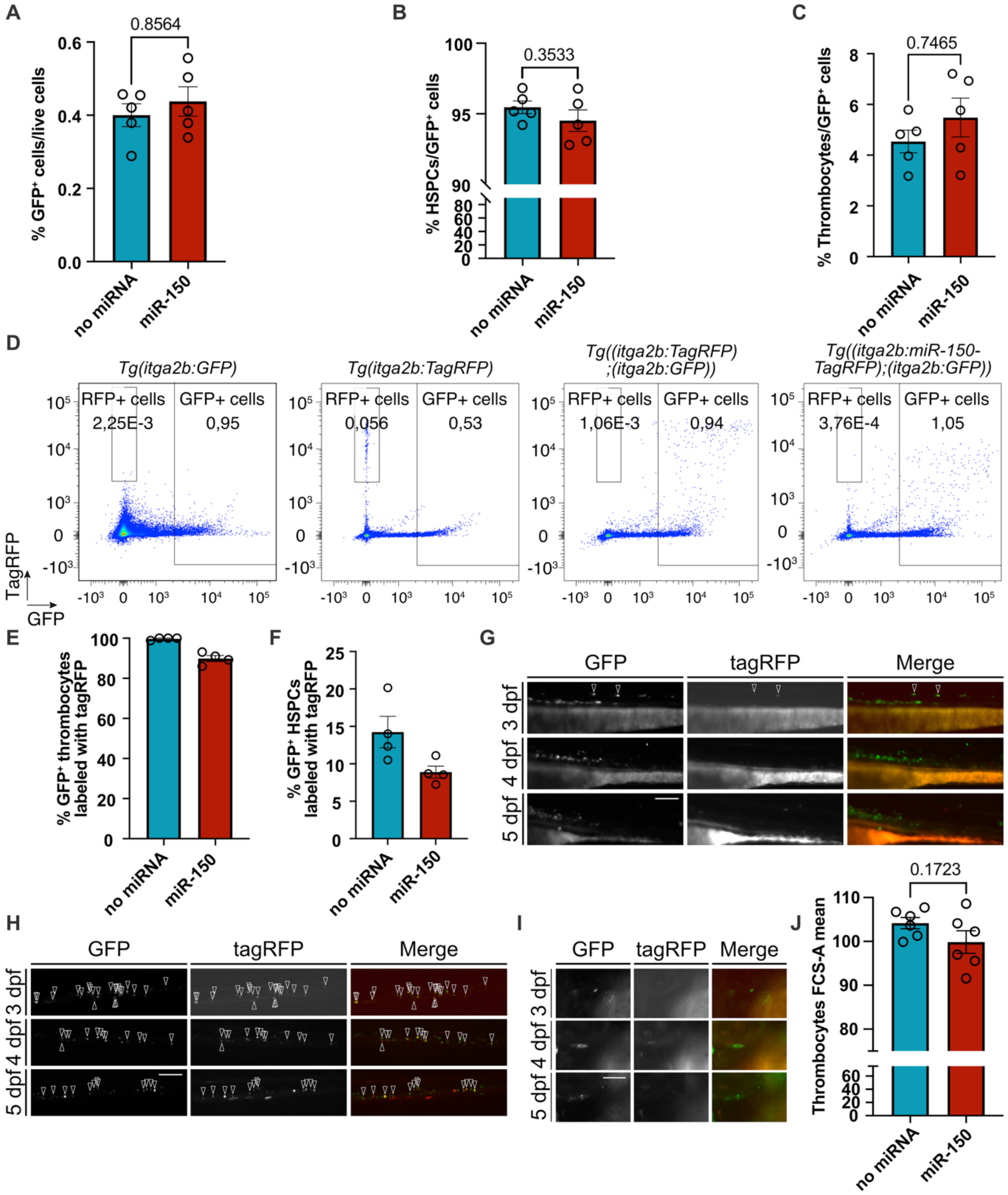


**Supplemental Figure 2. Effect of miR-150 upregulation on HSPC and thrombocyte differentiation, number and size.** (A) Percentage of the GFP⁺ cells in the whole population of living cells present in the double-transgenic zebrafish line (*Tg((itga2b:eGFP),(itga2b:miR-150-tagRFP))*) compared to the control line at 5 dpf (n = 5). (B) Percentage of the HSPCs in the GFP⁺ cell population in the double-transgenic zebrafish line (*Tg((itga2b:eGFP),(itga2b:miR-150-tagRFP))*) compared to the control line at 5 dpf (n = 5). (C) Percentage of thrombocytes in the GFP⁺ cell population in the double-transgenic zebrafish line (*Tg((itga2b:eGFP),(itga2b:miR-150-tagRFP))*) compared to the control line at 5 dpf (n = 5). (D) Examples of a flow cytometry plot (GFP – x-axis, tagRFP y-axis) used to distinguish between tagRFP⁺ cells and GFP⁺ cells, in the *Tg(itga2b:eGFP)*, *Tg(itga2b:tagRFP)*, *Tg((itga2b:eGFP),(itga2b:tagRFP))* and *Tg((itga2b:eGFP),(itga2b:miR-150-tagRFP))* transgenic lines at 5 dpf. (E) Percentage of tagRFP⁺ thrombocytes in the GFP⁺ thrombocytes population in the double-transgenic zebrafish line (*Tg((itga2b:eGFP),(itga2b:miR-150-tagRFP))*) compared to the control line at

5 dpf (n = 4). (F) Percentage of the tagRFP$^+$ HSPCs in the GFP$^+$ HSPCs population in the double-transgenic zebrafish line (*Tg((itga2b:eGFP),(itga2b:miR-150-tagRFP))*) compared to the control line at 5 dpf (n = 4). (G) *Tg(ita2b:eGFP)* transgene marks cells in the aorta/gonad/mesonephros (AGM) at 3-5 dpf, but *Tg(ita2b:miR-150-tagRFP)* does not. Arrow heads highlight only two cells that are both GFP$^+$/TagRFP$^+$ at 3 dpf. (H) Both *Tg(ita2b:eGFP)* and *Tg(ita2b:miR-150-tagRFP)* mark the same cells (arrow heads) in the caudal hematopoietic tissue (CHT) at 3-5 dpf. (I) *Tg(ita2b:eGFP)* transgene marks cells in the thymic lobes at 3-5 dpf, but *Tg(ita2b:miR-150-tagRFP)* does not. (J) The average size of the thrombocytes at 5 dpf in the double-transgenic zebrafish line (*Tg((itga2b:eGFP),(itga2b:miR-150-tagRFP))*) compared to the control line, assessed as a mean area of the forward scatter determined during a flow cytometry analysis of these cells (n = 6). Error bars represent SEM. Scale bar: 100 μm.

**Sup. Figure 3**

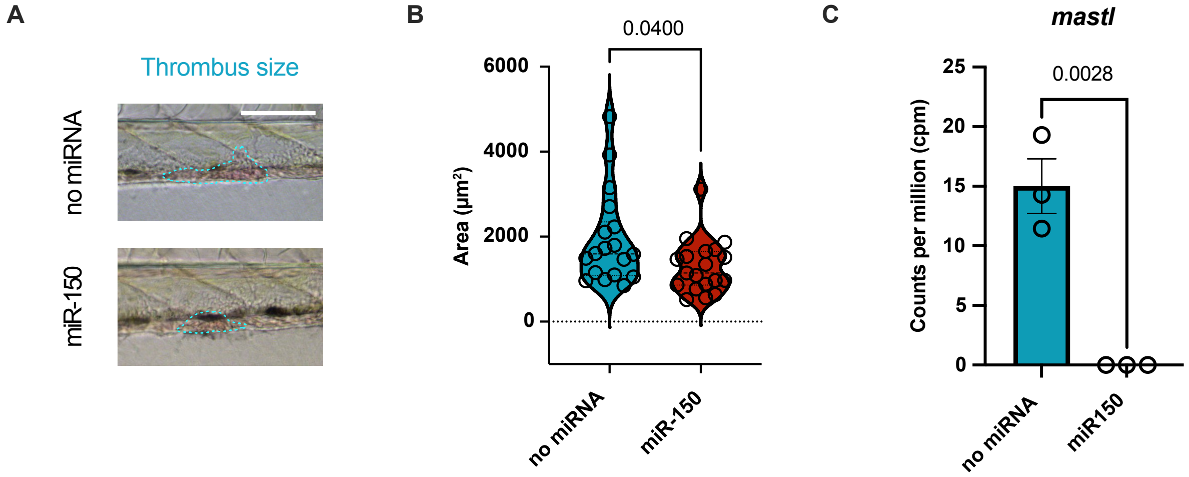


**Supplemental Figure 3. Thrombus formation after laser-induced vascular endothelial damage in control and miR-150-overexpressing fish.** (A) Representative bright-field images of a thrombus at 210 seconds after laser-induced caudal vein injury in the control and *Tg(itga2b:miR-150-tagRFP)* zebrafish larvae at 5 dpf. Scale bar: 100 µm. (B) Quantification of thrombus size at 210 seconds after the injury in the control - *Tg(itga2b:tagRFP)* (n = 18) and *Tg(itga2b:miR-150-tagRFP)* zebrafish (n = 19) at 5 dpf. (C) Quantification of the normalized counts of *mastl* transcript in the control: *Tg(itga2b:tagRFP)* and *Tg(itga2b:miR-150-tagRFP)* thrombocytes, obtain through the RNA sequencing analysis (n = 3). Error bars represent SEM.

Sup. Figure 4

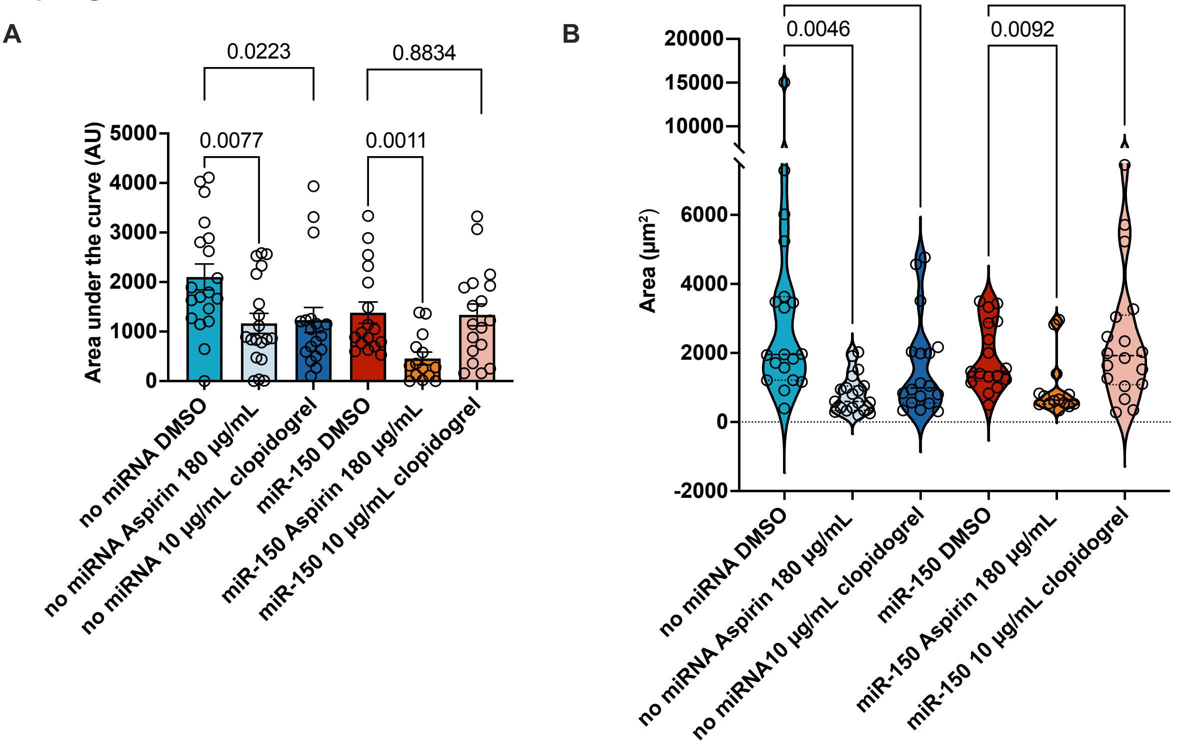


**Supplemental Figure 4. Effect of selected antiplatelet drug treatment on thrombus characteristics after laser injury in the miR-150-overexpressing line and controls.** (A) Quantification of area under the curve in graphs representing thrombocyte attachment at the site of the laser injury over time at 5 dpf in the control *Tg(itga2b:tagRFP)* or *Tg(itga2b:miR-150-tagRFP)* fish, treated with DMSO (n =19, 17), 180 µg/mL aspirin (n =19, 14) or 10 µg/mL clopidogrel (n = 18, 18, respectively) – as per graph **Figure 3A** and **3B**. Error bars represent SEM. (B) Quantification of thrombus size at 210 seconds after the injury at 5 dpf in the control - *Tg(itga2b:tagRFP)* and *Tg(itga2b:miR-150-tagRFP)* zebrafish treated with DMSO (n =19, 17), 180 µg/mL aspirin (n =19, 14) or 10 µg/mL clopidogrel (n = 18, 18).

**Sup. Figure 5**

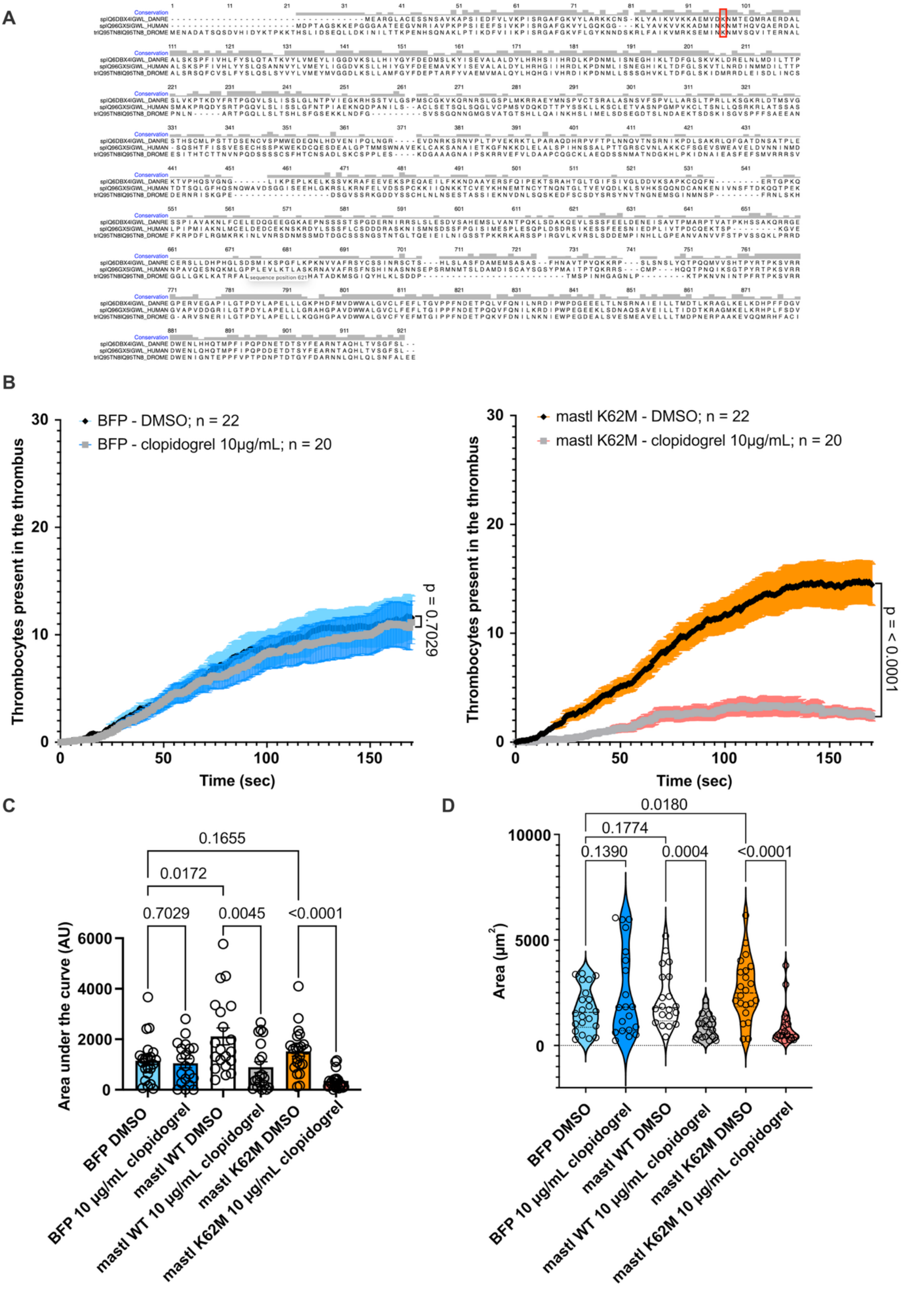


**Supplemental Figure 5. Effect of mastl overexpression in miR-150-overexpressing thrombocytes.** (A) The sequence of the *Drosophila* Greatwall protein (Q95TN8_ DROME) aligned with human MASTL (Q96GX5I_ HUMAN) and zebrafish mastl protein (Q6DB4I_ DANRE). Residues highlighted in red represent the hyperactive kinase mutation: *Drosophila* K97M, human K72M and zebrafish K62M. (B) Quantification of thrombocyte attachment at the site of laser injury over time at 5 dpf in *Tg(itga2b:miR-150-tagRFP)* fish, injected with Tol2-cryaa-Venus-itga2b-BFP (left) and cryaa-Venus-itga2b-mastl K62M mutant (right) construct and treated with DMSO (n = 22, 22) or 10 µg/mL clopidogrel (n = 20, 20, respectively). (C) Quantification of area under the curve in graphs representing the thrombocyte attachment at the site of the laser injury over time at 5 dpf in *Tg(itga2b:miR-150-tagRFP)* fish injected with BFP, mastl WT or mastl K62M mutant and treated with DMSO (n = 22, 20, 22) or 10 µg/mL clopidogrel (n = 20, 19, 20) – as per graph **Figure 3D** and **supplemental Figure 5B**. (D) Quantification of thrombus size at 210 seconds after the injury in *Tg(itga2b:miR-150-tagRFP)* fish injected with BFP, mastl WT or mastl K62M mutant and treated with DMSO (n = 22, 20, 22) or 10 µg/mL clopidogrel (n = 20, 19, 20) at 5 dpf. Error bars represent SEM.

Sup. Figure 6

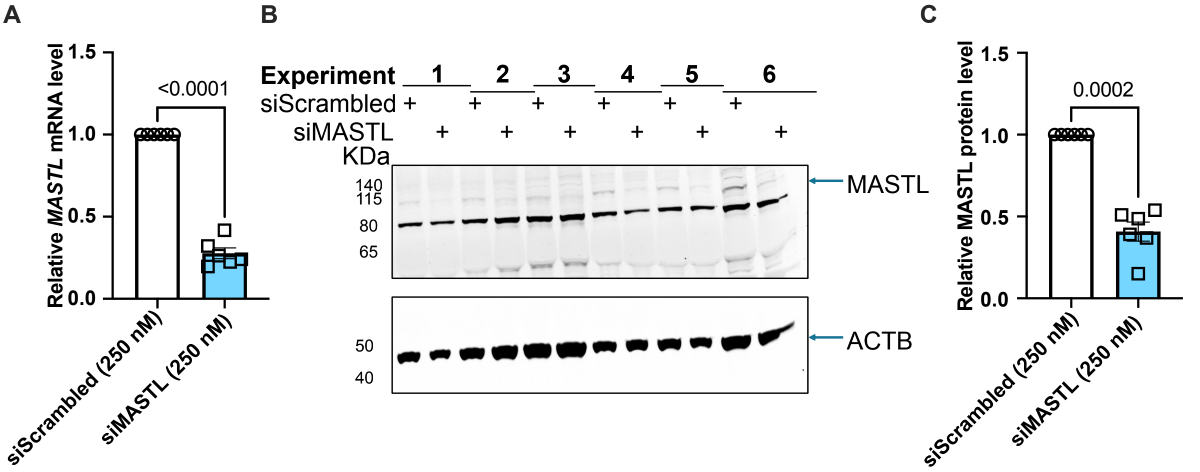


**Supplemental Figure 6. Expression of MASTL mRNA and protein in megakaryocytes nucleofected with MASTL-targeting siRNA.** (A) Expression of *MASTL* transcript in megakaryocytes, 48 hours after their nucleofection with scrambled siRNA (control) or *MASTL* targeting siRNA (siMASTL). Results are represented as a fold-change relative to control megakaryocytes (n = 6). (B) Western blot analysis of MASTL and ACTIN proteins in megakaryocytes, 48 hours after their nucleofection with scrambled siRNA (control) or *MASTL*-targeting siRNA (siMASTL). The blot shows six individual experiments. Actin was used for normalization. (C) Expression of MASTL protein in megakaryocytes, 48 hours after their nucleofection with scrambled siRNA (control) or *MASTL* targeting siRNA (siMASTL), as quantified using western blot analysis (**supplemental Figure 6B**). Results are represented as a fold-change relative to control megakaryocytes (n = 6). Error bars represent SEM.

### Methods

#### Ethical statement

Zebrafish (*danio rerio*) were raised and maintained according to FELASA guidelines.[1] All experiments, including the generation of transgenic animals were designed and performed in accordance with Swiss federal guidelines. Efforts were also made to comply to the 3R guidelines. No veterinary authorization was required as all experiments were performed on larvae on or before 5 days post-fertilization (dpf, before the animals were able to feed independently).

The human samples were obtained as part of an ADRIE study (Antiplatelet Drug Resistance and Ischemic Events; ClinicalTrials.gov identifier NCT00501423).[2,3] The study protocol was approved by the Central Ethics Committee of the University Hospitals of Geneva (Geneva center) and the Ethics Committee of Montpellier Saint-Eloi (Béziers and Montpellier centers). All patients included in the study signed a written informed consent.

#### Fish husbandry

Adult zebrafish (wild-type TU strain the transgenic strains; 3-24 months) were kept in a 14/10 h light/dark cycle, at 26 °C. We used the following transgenic animals: *Tg(-6.0itga2b:EGFP)*[4] and Tg(*kdrl:EGFP)*[s843].[5] Embryos obtained through natural matings were cultured at 28.5 °C in E3 medium. At 24-30 hours post-fertilization (hpf) the culture medium was supplemented with 0.003% (w/v) 1-phenyl 2-thiourea (PTU; Thermo Scientific Chemicals) to block the pigmentation and improve larval optical transparency.

#### Generation of transgenic animals

To generate each line plasmid DNA and *Tol2* mRNA were premixed and co-injected into one-cell-stage zygotes (20-22 pg of plasmid and 42 pg of mRNA). If needed, fish were raised to adulthood and further mated with wild-type TU zebrafish to identify founders with successful germline transmission and those yielding next generations with monoallelic expression were used to establish a transgenic line.

#### Whole-larvae genomic DNA isolation

16 anesthetized, wild-type TU larvae at 5 dpf were rinsed with E3 medium (1 mL) and water (1 mL). Liquid was removed using a drawn-out Pasteur pipette and DNA extraction buffer (160 µL) supplemented with Proteinase K (200 µg/mL, Roche) was added. The sample was incubated at 50 °C for about 3 h with occasional mixing. DNA was precipitated on ice by the addition of ethanol (Sigma-Aldrich) for 30 min. The sample was centrifuged at 12,000 rpm for 10 min at 4 °C. The supernatant was removed, and DNA was washed with 70% ethanol. The pelleted DNA was air-dried for 25 min and resuspended in water (100 µL; PanReac AppliChem).

#### Whole-embryo RNA isolation and cDNA synthesis

Wild-type TU zebrafish embryos 24 hpf (30 fish) were dechorionated and flash frozen in liquid nitrogen. TRIzol Reagent (500 uL; Invitrogen) was added and embryos were homogenized using a microfuge tube pestle and mortar. The homogenate was incubated at RT for 3 min, before the addition of chloroform (150 µL; Carlo Elba Reagents). The sample was vigorously shaken for 30 sec and incubated for 3 min at RT before centrifugation at 10'000 rpm for 15 min at 4 °C. The RNA-containing upper, aqueous phase was collected, and RNA was precipitated using 0.8 volume of 2-propanol (Sigma-Aldrich) at RT for 10 min. The sample was centrifuged at 10'000 rpm for 15 min at 4 °C and RNA was washed with 70% ethanol (Sigma-Aldrich). The pelleted RNA was air dried and resuspended in RNase-free water (50 µL; PanReac AppliChem). mRNA reverse transcription reactions were performed with the qScript cDNA SuperMix (Quanta bio).

#### Plasmid Generation

##### *Tol2-itga2b:TagRFP plasmid*

To generate the *Tol2-itga2b:TagRFP* construct, we first generated a *Tol2-itga2b:miR-223-TagRFP* plasmid. The *miR-223-tagRFP*-coding region upstream of a Kozak sequence was PCR amplified from the *Tol2-lyzC-RFP-miR-223* plasmid (Qing Deng, Addgene)[6] using the following primers: 5′-aagcttgatatcgaattcctgcagcccgggccaccatggatgaggaaatc-3′ and 5′-cagatctagcgctccgccgatttcggcctattg-3′. *Tol2-itga2b:dsRED-pri-miR-223* plasmid (previously prepared in our lab by Dr. Veronika Zapilko) was digested with XmaI and ZraI. A small fragment of a *Tol2 arm* lost during the digestion process was PCR amplified using the following primers: 5'-cgaaatcggcggagcgctagatctgcgaagatacggcc-3' and 5'-tgttcattggtcctttggaagtgacgtcatgtcacatctattaccacaatg-3'. The *Tol2- itga2b:miR223-TagRFP* plasmid was assembled from the digested vector and the two PCR products (*miR-223-tagRFP* and *Tol2 arm*) using HiFi DNA Assembly Master Mix (NEB). The plasmid was sequenced to confirm the successful ligation.

Since the *miR-223-tagRFP* coding sequence, generated from the *Tol2-lyzC-RFP-miR-223* plasmid, contained the first exon (including ATG) and first intron of *actb1*, with the original *pri-miR-30* miRNA sequence backbone and pre-miR-223 sequence both inserted into the intron,[6,7] we decided to remove the *pre-miR-30* sequence backbone. To do so the *5'-actb1* and *3'-actb1*-coding regions were PCR amplified from the *Tol2- itga2b:miR223-TagRFP* plasmid using the following primers containing MluI site (underlined): *5'-actb1* (5'-atcgaattcctgcagcccgggccaccatggatgaggaaatcgctg-3' and 5'-tttattagatctacgcgtgactttcaaagctg-3') and *3'-actb1* (5'-gaaagtcacgcgtagatctaataaaagttaatttaagcgttcattacaataaattcaagtgaatctg-3' and 5'-taatcagctcttcgcccttagacaccatggtggcgaccggtacccccg-3'). The MluI site was introduced to allow a simple introduction of any desired pri-miRNA sequence. *Tol2-itga2b:miR223-TagRFP* plasmid was digested with NcoI-HF and the *Tol2-itga2b:TagRFP* was assembled from the resulting vector and the *5'-actb1* and *3'-actb1* PCR products using HiFi DNA Assembly Master Mix (NEB). The plasmid was sequenced to confirm the successful ligation.

#### *Tol2-itga2b:miR150-TagRFP plasmid*

To generate the *Tol2- itga2b:miR-150-TagRFP* plasmid, the pre-miR-150-coding region was PCR amplified from whole larvae genomic DNA using the following primers containing MluI site (underlined): 5′- cgaagttcttcagctttgaaagtcacgcgttggactttccacggtaatgtttg-3′ and 5′-cttaaattaacttttattagatctacgcgtgactaaatggacatgaggg-3′. *Tol2-itga2b:TagRFP* plasmid was digested with MluI-HF and the *Tol2-itga2b:miR-150-TagRFP* was assembled from the resulting vector and the *pre-miR-150* PCR products using HiFi DNA Assembly Master Mix (NEB). The plasmid was sequenced to confirm the successful ligation.

#### *Tol2-cryaa:Venus-itga2b:mastl WT plasmid*

To generate the *Tol2-cryaa:Venus-itga2b:mastl WT* construct, first we generated a *Tol2-cryaa:Venus-itga2b:mCherry* plasmid. The *cryaa:Venus*-coding region was PCR amplified from *pKE4-I-SceI-fabp10a:Xpt-β-cat, cryaa:Venus* plasmid (Kimberley Evason, Didier Stainier, Addgene)[8] using the following primers: 5′-actcactatagggcgaattgggtacctccccagcatgcctgctattg-3′ and 5′-ttactagtggatccgagctcggtacattaatagtgtgcattcagtgcagg-3′. *Tol2-itga2b:mCherry* plasmid (previously prepared in our lab) was digested with KpnI-HF, and the *Tol2-cryaa:Venus-itga2b:mCherry* was assembled from the resulting vector and the *cryaa:Venus* PCR products using HiFi DNA Assembly Master Mix (NEB). The plasmid was sequenced to confirm the successful ligation.

Next, zebrafish *mastl*-coding sequence was PCR amplified from 24 hpf embryo cDNA using the following primers: 5'-tcgataagcttgatatcgaattcctgcagcccgggccaccatggaagctcgtggattgg-3' and 5'-atggtacagtaaaacgacggccaggatccaccggttagagactgaagccggaaacggtc-3'. The *Tol2-cryaa:Venus-itga2b:mCherry* plasmid was digested with AgeI-HF and XmaI and the *Tol2-cryaa:Venus-itga2b:mastl WT* was assembled from the resulting vector and the *mastl* PCR product using HiFi DNA Assembly Master Mix (NEB). The plasmid was sequenced to confirm the successful ligation.

#### *Tol2-cryaa:Venus-itga2b:mastl K62M plasmid*

To generate the *Tol2-cryaa:Venus-itga2b:mastl K62M* construct, the *mastl K62M*-coding region was PCR amplified from *Tol2-cryaa:Venus-itga2b:mastl WT* plasmid in two fragments, both containing the K62M mutation in the overlapping parts of the amplicons. The PCR fragments were prepared using the following primers: *mastl fragment 1* (5′-tcgataagcttgatatcgaattcctgcagcccgggccaccatggaagctcgtggattgg-3′ and 5′-gtcatgttcatatccaccatttctgctttc-3′), *mastl fragment 2* (5'-tggtggatatgaacatgactgagcagatga-3' and 5'-atggtacagtaaaacgacggccaggatccaccggttagagactgaagccggaaacggtc-3'). The *Tol2-cryaa:Venus-itga2b:mCherry* plasmid was digested with AgeI-HF and XmaI and the *Tol2-cryaa:Venus-itga2b:mastl K62M* was assembled from the resulting vector and the two *mastl K62M* PCR products using HiFi DNA Assembly Master Mix (NEB). The plasmid was sequenced to confirm the successful ligation.

#### *Tol2-cryaa:Venus-itga2b:BFP plasmid*

To generate the *Tol2-cryaa:Venus-itga2b:BFP* construct, the *BFP*-coding region was PCR amplified from *Tol2-BFP* plasmid (gift from Julien Bertrand) using the following primers: 5′-tgatatcgaattcctgcagcccgggccaccatggaagcagcaagagcaagccaaaggtgagcaagggcgaggagct-3′ and 5′-

aaaacgacggccaggatccaccggttacttgtacagctcgtccatgcc-3'. The *Tol2-cryaa:Venus-itga2b:mCherry* plasmid was digested with AgeI-HF and XmaI and the *Tol2-cryaa:Venus-itga2b:mastl K62M* was assembled from the resulting vector and the *BFP* PCR product using HiFi DNA Assembly Master Mix (NEB). The plasmid was sequenced to confirm the successful ligation.

### Cell sorting and flow cytometry

#### *Larvae dissociation into single cells*

Anesthetized larvae at 5 dpf and were incubated with 0.5 mg/mL Liberase TM (Thermolysin Medium; Roche) solution in DPBS (Gibco) for 2 x 45 min at 33 °C with agitation. After 45 min and 90 min tissues were disaggregated by pipetting. At 90 min the digestion was blocked by addition of equal volume of 1% fetal calf serum/1 mM EDTA (Invitrogen) in 0.9 x PBS. Cell suspension was centrifuged at 1000 rpm, for 10 min at 4 °C, supernatant was removed, and cells were resuspended in 1% fetal calf serum/1 mM EDTA in 0.9 x PBS solution. Cells were passed through 70 µm cell strainer prior to flow cytometry. Dead cells were excluded by staining with DAPI (Thermo Fischer Scientific). Cell sorting was performed using an Aria II (BD Biosciences, software diva v6.1.3) sorter and all data were analyzed with FlowJo 10. Cells were sorted into RLT buffer (Qiagene) and kept on ice before RNA extraction.

#### *Quantification of thrombocytes/HSPCs number and size*

*Tg(itga2b:miR-150-tagRFP)* or *Tg(itga2b:tagRFP)* fish were crossed to *Tg(itga2b:EGFP)* animals and the larvae were screened at 4 dpf for the presence of both transgene ($GFP^+$/$TagRFP^+$ cells). At 5 dpf 30+ larvae per experimental condition were collected and dissociated into single cells (as described above). The analysis was performed using an Aria II (BD Biosciences, software diva v6.1.3) sorter. At least 100'000 live cell events were collected for the analysis. The distinction of the HSPCs was made based on the presence of $GFP^{low}$ signal (low GFP intensity) and larger forward scatter area (FSC-A). The thrombocytes were characterized by the presence of $GFP^{high}$ signal (high GFP intensity) and small FCS-A size. For the thrombocytes size measurement, the average FCS-A of all thrombocytes present in the sample was calculated.

#### *Isolation of vascular endothelial cells*

*Tg(itga2b:miR-150-tagRFP)* or *Tg(itga2b:tagRFP)* fish were crossed to *Tg(kdrl:EGFP)* animals and the larvae were screened at 4 dpf for the presence of both transgene ($GFP^+$/$TagRFP^+$ cells). At 5 dpf, no less than 30 larvae per experimental condition were collected and dissociated into single cells (as described above).

### Laser-induced vascular endothelial injury and data analysis

#### *Laser set-up*

To perform the injury, laser was focused on the endothelial wall of the posterior cardinal vein within the fifth somite caudal to the anal pore. The laser was set to the power, aperture and speed of 20, 15 and 20, respectively.

#### *Exclusion criteria*

Fish that twitched during the laser injury or fish that were not optimally mounted (leading to blurry images) were excluded from the analyses.

#### *Laser injury analysis*

To analyze the number of thrombocytes in the thrombus at a given timepoint (frame), we manually counted the fluorescent thrombocytes present at the site of injury, by adding the thrombocytes that aggregated and subtracting those that left the wound. The thrombocyte was treated as part of a thrombus if its localization within the thrombus was consistent over at least 3 frames. The analysis was performed using the imaging analysis software Fiji and the number of thrombocytes present in the thrombus was plotted against time.

To calculate the area of a thrombus (2D) brightfield images of the wound were utilized. The shape of the thrombus was manually drawn using the imaging analysis software Fiji, the size of the thrombus was calculated and reported in $\mu m^2$.

### RNA extraction

Complete RNA, containing both mRNAs and miRNAs, was isolated from sorted thrombocytes, vascular endothelial cells or *in vitro* generated megakaryocytes (MKs) using the miRNeasy Tissue/Cells Advanced Mini Kit (Qiagen) following the manufacturer's protocol.

### Reverse transcription-qPCR

#### *miRNA quantitative real-time PCR*

The appropriate miRNAs were reversed transcribed individually using the TaqMan$^{TM}$ MicroRNA Reverse Transcription Kit (Applied Biosystems). The commercial, predesigned probes (RT, Applied Biosystems) were utilized for dre-miR-150 miR-126, miR-223 and U6 snRNA (an internal control for normalization). The catalog numbers of all the probes are provided in **supplementary Table S4**. The quantification of each sample was performed using individual, commercially available, predesigned TagMan$^{TM}$ probes (TM, Applied Biosystems) and TaqMan$^{TM}$ Fast Advanced Master Mix (Applied Biosystems).

#### *mRNA quantitative real-time PCR*

The mRNAs reverse transcription reaction was performed with the ImProm-II$^{TM}$ Reverse Transcription System (Promega). The quantitative real-time PCR (qPCR) was achieved using PowerUp$^{TM}$ SYBR$^{TM}$ Green Master Mix (Applied Biosystems). Human β2-microglobulin (HB2M) was used as internal control for mRNA normalization. All primers are listed in **supplementary Table S4.**

#### *qPCR run and analysis*

The qPCRs runs were performed using a QuantStudio 1 Real-Time PCR System and analyzed using QuantStudio Design & Analysis Software v1.5.2. (Thermofisher). The relative expression level of transcripts was evaluated using the $2^{-\Delta\Delta Ct}$ formula. Each qPCR sample was quantified in triplicates and averages of the triplicates from a single sample were used. Each experiment was repeated at least three times, and fold-change averages of all experiments were combined.

### RNA sequencing

#### *Library preparation, sequencing, read mapping to the reference genome and gene coverage reporting*

Following thrombocyte sorting and RNA isolation, the quality of each sample was assessed using the Agilent 2100 Bioanalyzer with the Agilent RNA 6000 Nano Kit (Agilent Technologies). cDNA libraries were constructed using the SMARTer Ultra Low Input RNA kit (Clontech), combined with Nextera XT DNA Library preparation kit (Illumina) according to the manufacturers' protocols. Libraries were sequenced using single-end reads (100 nt-long) on an Illumina NovaSeq 6000. The sequencing was performed twice to achieve the appropriate read depth. Fastq reads were mapped to the Ensembl *Danio rerio* GRCz11 genome with STAR v.2.7.10b.[9] The quality analysis was performed with the Picard tool (http://broadinstitute.github.io/picard/). Quantification of transcripts expression was performed with HTSeq v0.11.3 (htseq-count).[10] Sequence data have been submitted to the GEO repository (accession number: GSE307726).

#### *RNAseq analysis*

After normalization, the poorly detected genes were filtered. Out of the 32'520 total genes in the Ensembl annotation, 18'990 genes with a count above 10 were kept for the further analysis. The differential expression analysis was performed with the statistical analysis R/Bioconductor package edgeR 1.34.1..[11] Only genes with a multiple testing Benjamini and Hochberg correction FDR 5% and a fold change threshold of 2 were considered. The volcano plot was generated using VolcaNoseR web app (https://huygens.science.uva.nl/VolcaNoseR/).[12]

### Drug exposure and analysis

At 4.5 dpf larvae were placed overnight in E3 medium supplemented with either 180 µg/mL O-Acetylsalicylic acid (aspirin; Thermo Scientific Chemicals) or 10 µg/mL (S)-(+)-Clopidogrel hydrogensulfate (clopidogrel; Sigma-Aldrich). Larvae treated with an equivalent amount of DMSO (Sigma-Aldrich) vehicle were used as controls. To identify a possible effect of these drugs on thrombus formation, at 5 dpf we analyzed laser-induced thrombocyte aggregation over time and evaluated thrombus size as described in Methods.

### *In vitro* generation of platelets derived from human hematopoietic stem cells

The purification and differentiation of human hematopoietic stem cells ($CD34^+$) were performed as previously described.[13] In brief, the $CD34^+$ cells were obtained from the buffy coats of healthy adult volunteers from the Geneva University Hospitals blood bank (Hôpitaux universitaires de Genève, HUG). The cells were isolated using a CD34 MicroBead Kit (Miltenyi Biotec). Cells were seeded at 4 x $10^4$ cells/mL and cultured for 7 days in StemSpan Serum-Free Expansion Medium (Stemcell) containing 20 ng/ml human low-density lipoprotein (LDL, Stemcell technologies), 1 µM StemRegenin 1 (SR1, Cellagen Technology), 1x StemSpan™ Megakaryocyte Expansion Supplement (Stemcell Technologies) and 1x penicillin-streptomycin-glutamine (Gibco). Megakaryocyte (MK) progenitors were resuspended at 5 x $10^4$ cells/mL and cultured in the presence of 0.5 µg/mL of thrombopoietin (TPO, Miltenyi Biotec) for 8 additional days to allow their differentiation into MKs and platelets.

#### Nucleofection procedure and downregulation of MASTL

$CD34^+$ cells were resuspended at 5 x $10^5$ cells/mL in StemSpan Serum-Free Expansion Medium containing LDL and SR1 on day 12. The next day cells were centrifuged at 1600 rpm for 10 minutes at RT. Subsequently, 1.5 x $10^6$ cells per experimental condition were resuspended in a solution containing siRNAs (250 nM; Qiagen) and P3 primary cell solution 4D-Nucleofector™ X Kit S (Lonza). Cells transfected using 4D-Nucleofector X Unit (Lonza), program DZ-100. After a 10 min rest period, cells were seeded at 5 x $10^5$ cells/mL in a 24-well plate with 2 mL of prewarmed media (Stemspan) supplemented with LDL, SR1, and 0.5 µg/mL TPO. All subsequent assays were performed on day 15. The sequences of the siRNAs used is provided in **supplementary Table S4,** a scrambled siRNA was used as a negative control.

#### Western blot

A Western blot analysis, to quantify the MASTL protein, was performed 48 hours after nucleofection. The quantification was performed in megakaryocytes. Proteins were labelled with an anti-MASTL rabbit polyclonal antibody (dilution 1:500, Abcepta) and an anti-β-actin mouse monoclonal antibody (dilution 1:12'000, Sigma-Aldrich). Goat anti-rabbit Dylight 680 secondary antibody (dilution 1:5000, Rockland) and goat anti-mouse Dylight 800 secondary antibody (dilution 1:10000, Rockland) were used for detection of a fluorescent signal in an Odyssey M imager (LicorBio). Band intensities were quantified using the Fiji software (http//imagej.nih.gov/ij), and the MASTL protein level was normalized to β-actin.

#### VASP assay in human platelets

Platelets, obtained by *in vitro* differentiation of megakaryocytes (described above), were resuspended in approximately 120 µL of Tyrode's albumin buffer. Subsequently, 40 µL of platelets suspension was incubated for 5 min in the presence of 50 nM cangrelor tetrasodium salt (cangrelor; Sigma-Aldrich) or water as a negative control. Immediately, after the treatment, the VASP phosphorylation status and the platelet reactivity index (PRI) of these cells were evaluated using a standardized assay (Platelet VASP/P2Y12; Biocytex), according to the manufacturer's instructions.[14] The analysis was performed using an Accuri C6 (BD Biosciences, software CFlow Plus v1.0.264.15) flow cytometer. At lease 5'000 platelets were collected for the analysis. All data were analyzed with FlowJo 10.

#### Cardiovascular Patients blood collection

Blood was collected by venipuncture to ethylenediaminetetraacetic acid (EDTA)-containing tubes. Plasma was acquired by a single centrifugation of whole blood at 2300 × *g* for 15 min at RT and was stored at −80 °C until analysis.

#### Quantification of microRNAs in plasma obtained from cardiovascular patients

After thawing, 200 µL of plasma samples were centrifuged at 1000 × *g* for 5 min at 4 °C. The *Caenorhabditis elegans* cel-miR-39 (5 fmol, Ambion) was added to each sample as a spike-in control to assess the efficiency of the purification and reverse transcription procedures. miRNA was isolated using miRNeasy Serum/Plasma Advanced Kit (Qiagen) following the manufacturer's instructions. A fixed volume of RNA (2 µL) was utilized in the reverse transcription reaction with preamplification performed using TaqMan Advanced miRNA cDNA Synthesis Kit (Applied Biosystem). The quantification of each sample was performed in triplicates using TaqMan™ Fast Advanced Master Mix (Applied Biosystems) and the individual, commercially available, predesigned probes (Applied Biosystems). The catalog numbers of the probes are provided in **supplementary Table S4**. The qPCRs run was performed using a 7900HT SDS system (Applied Biosystems). Firstly, to determine, whether the RNA extraction procedure was consistent between all samples, we analyzed the level of cel-miR-39, spiked-in before the extraction was performed, by qPCR and showed a stable mean value of

18.84±11.15 % Ct. The previously described, three stably expressed, endogenous miRNAs: hsa-miR-16-5p, hsa-miR-93-5p and hsa-miR-484, were used as a reference for normalization in our study.[15] To confirm their suitability we determined the geNorm values for these miRNAs in our sample set to be below the cut-off 1.5 (specifically: 0.5019, 0.5839 and 0.5898 for miR-16-5p, hsa-miR-93-5p and hsa-miR-484, respectively).[16,17] The relative level of each miRNA was determined according to the previously described protocol.[17] Exclusion: For hsa-miR-142-5p two patient pairs were excluded, due to undetectable miR-142-5p in one of the patients in each pair.

### Statistical Analysis

Data presented in this publication are shown as a mean and a standard error of the mean (SEM) of at least three independent experiments. The unpaired/paired student's t-test or the one-way analysis of variance (ANOVA) followed by a posthoc Tukey test were performed, when appropriate. A *p*-value $< 0.05$ was considered as significant. All graphs were generated using GraphPad Prism10 software (v10.2.2, Dotmatics).

***Supplemental Table S1*** *Extreme phenotype patients in the cohort.*

| | LPR: VASP<16 | HPR: VASP>50 |
|---|---|---|
| **n** | 17 | 17 |
| **VASP-PRI (% mean ± SD)** | 9.1±3.9 | 62.8±9.4 |
| **CYP2C19 het mutant (%)** | 5.9% | 5.9% |
| **Age (mean ± SD)** | 62.7±9.4 | 62.5±8.5 |
| **Weight (kg; mean ± SD)** | 73.2±9.7 | 72.8±10.1 |
| **Diabetes (%)** | 23.5% | 23.5% |
| **Female (%)** | 29.4% | 23.5% |

***Supplemental Table S2*** *List of significantly up and downregulated transcripts in miR-150-overexpressing thrombocytes. Mastl transcript is highlighted in red.*

| ENSEMBL | SYMBOL | logFC | logCPM | LR | PValue | FDR |
|---|---|---|---|---|---|---|
| ENSDARG00000056750 | NA | -13.061 | 4.047 | 37.804 | 7.82324446579813e-10 | 1.48563412405506e-05 |
| ENSDARG00000020028 | cps1 | -13.503 | 4.486 | 34.234 | 4.88589994470022e-09 | 4.63916199749286e-05 |
| ENSDARG00000007655 | crybb1l3 | -14.282 | 5.273 | 28.719 | 8.36816652893218e-08 | 0.0005 |
| ENSDARG00000077193 | nags | -13.105 | 4.090 | 28.010 | 1.20684746002228e-07 | 0.0006 |
| ENSDARG00000007576 | crybb1l1 | -7.908 | 12.716 | 25.217 | 5.122462694581e-07 | 0.0017 |
| ENSDARG00000041925 | cryba2b | -8.365 | 11.082 | 24.983 | 5.78238472459015e-07 | 0.0017 |
| ENSDARG00000058133 | foxd2 | -11.982 | 3.011 | 24.731 | 6.59062547852589e-07 | 0.0017 |
| ENSDARG00000014803 | cryba1l2 | -7.665 | 7.233 | 24.352 | 8.02579359126381e-07 | 0.0017 |
| ENSDARG00000030349 | cryba2a | -7.255 | 9.962 | 24.198 | 8.69322689728726e-07 | 0.0017 |
| ENSDARG00000053862 | crygmx | -8.178 | 12.017 | 23.989 | 9.69080624561582e-07 | 0.0017 |
| ENSDARG00000021889 | gja3 | -12.821 | 3.808 | 23.759 | 1.09206047131266e-06 | 0.0017 |
| ENSDARG00000016793 | crybb1l2 | -8.261 | 10.015 | 23.675 | 1.14084027786762e-06 | 0.0017 |
| ENSDARG00000032929 | cryba1l1 | -7.873 | 12.155 | 23.598 | 1.18725201241375e-06 | 0.0017 |
| **ENSDARG00000055566** | **mastl** | **11.058** | **3.073** | **23.357** | **1.34570473106676e-06** | **0.0017** |
| ENSDARG00000030411 | crygn2 | -7.739 | 11.156 | 23.227 | 1.43977185632804e-06 | 0.0017 |
| ENSDARG00000062147 | otc | -4.615 | 6.534 | 23.157 | 1.49265335724246e-06 | 0.0017 |
| ENSDARG00000024548 | cryba4 | -7.825 | 11.933 | 23.155 | 1.49474046536767e-06 | 0.0017 |
| ENSDARG00000053875 | cryba1b | -7.668 | 12.232 | 22.385 | 2.23066420495509e-06 | 0.0023 |
| ENSDARG00000068507 | crybb1 | -7.954 | 12.317 | 22.329 | 2.29657914224737e-06 | 0.0023 |
| ENSDARG00000099579 | aldh18a1 | -5.211 | 7.410 | 22.121 | 2.56018386192707e-06 | 0.0024 |
| ENSDARG00000045898 | NA | -5.491 | 7.856 | 21.656 | 3.26201858484432e-06 | 0.0029 |
| ENSDARG00000017299 | fabp4a | -6.271 | 8.247 | 21.021 | 4.54351771163224e-06 | 0.0036 |
| ENSDARG00000042470 | s1pr3a | -12.020 | 3.034 | 21.016 | 4.55433823906779e-06 | 0.0036 |
| ENSDARG00000091692 | havcr2 | -12.278 | 3.270 | 21.001 | 4.59082155350783e-06 | 0.0036 |
| ENSDARG00000017036 | svilc | -12.612 | 3.630 | 20.598 | 5.66616893726292e-06 | 0.0043 |
| ENSDARG00000104801 | tubb6 | -15.218 | 6.202 | 20.393 | 6.30497435905556e-06 | 0.0046 |
| ENSDARG00000008131 | sox1b | -9.806 | 4.794 | 20.079 | 7.43141406873354e-06 | 0.0052 |
| ENSDARG00000074919 | NA | -7.592 | 5.807 | 19.738 | 8.88345795481228e-06 | 0.0060 |
| ENSDARG00000053323 | zgc:112285 | -12.520 | 3.521 | 19.259 | 1.14149062387165e-05 | 0.0075 |
| ENSDARG00000094310 | si:ch211-255g12.6 | -7.267 | 9.673 | 18.925 | 1.3598667009861e-05 | 0.0086 |
| ENSDARG00000042641 | cyp51 | -3.551 | 5.020 | 18.363 | 1.82589456832041e-05 | 0.0112 |
| ENSDARG00000039522 | tubb2 | -11.594 | 2.639 | 18.249 | 1.93791178809596e-05 | 0.0114 |
| ENSDARG00000091633 | si:dkey-177p2.18 | -11.366 | 2.373 | 18.209 | 1.97919170548469e-05 | 0.0114 |
| ENSDARG00000041952 | prox2 | -8.810 | 5.819 | 18.029 | 2.17552593989095e-05 | 0.0122 |
| ENSDARG00000015445 | lim2.4 | -9.015 | 9.377 | 17.758 | 2.50809394782555e-05 | 0.0136 |
| ENSDARG00000116164 | crygm2d8 | -6.380 | 8.919 | 17.529 | 2.82912406212608e-05 | 0.0149 |
| ENSDARG00000071139 | LOC100151335 | -13.392 | 4.375 | 17.343 | 3.11968377236603e-05 | 0.0160 |
| ENSDARG00000076572 | crygm2d7 | -6.604 | 7.547 | 17.042 | 3.65690866506299e-05 | 0.0173 |

| | | | | | | |
|---|---|---|---|---|---|---|
| ENSDARG00000036834 | krt92 | -4.637 | 7.246 | 17.014 | 3.70968150581023e-05 | 0.0173 |
| ENSDARG00000041848 | rh50 | -10.290 | 1.685 | 17.013 | 3.71319049790319e-05 | 0.0173 |
| ENSDARG00000109861 | crygm2d19 | -5.959 | 8.763 | 16.998 | 3.74151919144746e-05 | 0.0173 |
| ENSDARG00000077461 | dhx32a | -13.286 | 4.414 | 16.791 | 4.17396642636838e-05 | 0.0189 |
| ENSDARG00000057460 | crygm2d13 | -6.209 | 8.932 | 16.674 | 4.43849870051928e-05 | 0.0196 |
| ENSDARG00000097003 | NA | -11.040 | 2.055 | 16.574 | 4.67987540982104e-05 | 0.0202 |
| ENSDARG00000013963 | mipb | -9.947 | 10.308 | 16.399 | 5.13106194418269e-05 | 0.0217 |
| ENSDARG00000069827 | crygm2d11 | -6.172 | 6.475 | 16.200 | 5.6985914446654e-05 | 0.0235 |
| ENSDARG00000079305 | hbae3 | -5.067 | 5.748 | 16.063 | 6.12620363040418e-05 | 0.0248 |
| ENSDARG00000079496 | bicd1a | -10.618 | 1.645 | 15.774 | 7.13874445392499e-05 | 0.0282 |
| ENSDARG00000087324 | crygm2d1 | -5.863 | 8.170 | 15.561 | 7.99028205147109e-05 | 0.0310 |
| ENSDARG00000090170 | rab11fip4a | -11.607 | 2.985 | 15.313 | 9.10726877699241e-05 | 0.0346 |
| ENSDARG00000090997 | vegfba | -11.025 | 2.307 | 15.036 | 0.0001 | 0.0385 |
| ENSDARG00000040245 | kpnb3 | -3.391 | 6.705 | 15.013 | 0.0001 | 0.0385 |
| ENSDARG00000037402 | lim2.3 | -11.050 | 7.800 | 15.002 | 0.0001 | 0.0385 |
| ENSDARG00000001463 | tdh2 | -6.259 | 4.492 | 14.628 | 0.0001 | 0.0460 |
| ENSDARG00000088823 | crygm2d3 | -5.555 | 7.764 | 14.580 | 0.0001 | 0.0464 |
| ENSDARG00000101959 | etv1 | 9.730 | 2.118 | 14.456 | 0.0001 | 0.0486 |
| ENSDARG00000089429 | si:dkey-205h13.2 | 12.330 | 3.494 | 14.389 | 0.0001 | 0.0491 |
| ENSDARG00000042189 | tspan33b | -12.756 | 3.751 | 14.373 | 0.0001 | 0.0491 |

***Supplemental Table S3*** *List of Reagents*

| **Reagent** | **Source** | **Identifier** |
|---|---|---|
| **Antibodies** | | |
| Rabbit anti-MASTL | Abcepta | Cat# AP7147d |
| Mouse anti-β-actin | Sigma-Aldrich | Cat# A5441 |
| Goat anti Rabbbit IgG DyLight 680 Conjugated Antibody | Rockland | Cat# 611-144-122 |
| Goat anti Mouse IgG DyLight 800 Conjugated Antibody | Rockland | Cat# 610-145-121 |
| **Chemicals** | | |
| (S)-(+)-Clopidogrel hydrogensulfate (clopidogrel) | Sigma-Aldrich | Cat# SML0004 |
| 1-phenyl 2-thiourea (PTU) | Thermo Scientific Chemicals | Cat# 207250050 |
| 2-Propanol | Sigma-Aldrich | Cat# 59300 |
| Agarose, low gelling temperature | Sigma-Aldrich | Cat# A9045 |
| Cangrelor tetrasodium salt | Sigma-Aldrich | Cat# SML2004 |
| Chloroform | Carlo Erba Reagents | Cat# P02410E16 |
| DAPI | Invitrogen | Cat# D1306 |
| Dimethyl sulfoxide (DMSO) | Sigma-Aldrich | Cat# D8418 |
| DPBS, no calcium, no magnesium | Gibco | Cat# 14190 |
| Ethyl 3-aminobenzoate methanesulfonate (MS-222) | Sigma-Aldrich | Cat# E10521 |
| Ethyl alcohol, pure (ethanol) | Sigma-Aldrich | Cat# 51976 |
| human LDL | Stemcell technologies | Cat# 02698 |
| Human TPO | Miltenyi Biotec | Cat# 130-095-754 |
| Liberase TM Research Grade | Roche | Cat# 05401119001 |
| O-Acetylsalicylic acid (aspirin) | Thermo Scientific Chemicals | Cat# A12488 |
| penicillin-streptomycin-glutamine | Gibco | Cat# 10378016 |
| Protinase K, recombinant, PCR Grade | Roche | Cat# 03115887001 |
| StemRegenin 1 (SR1) | Cellagen Technology | Cat# C7710-1s |
| StemSpan Serum-Free Expansion Medium | Stemcell technologies | Cat# 09650 |
| StemSpan™ Megakaryocyte Expansion Supplement (CC220) | Stemcell technologies | Cat# 02696 |
| TRIzol Reagenet | Invitrogen | Cat# 15596018 |
| UltraPure 0.5M EDTA, pH 8.0 | Invitrogen | Cat# 15575020 |
| Water for molecular biology | PanReac AppliChem ITW Reagents | Cat# A7398 |
| **Kits** | | |
| CD34 MicroBead Kit, human | Miltenyi Biotec | Cat# 130-100-453 |
| HiFi DNA Assembly Master Mix | NEB | Cat# E2621 |
| miRNeasy Serum/Plasma Advanced Kit | Qiagen | Cat# 217204 |
| miRNeasy Tissue/Cells Advanced Mini Kit | Qiagen | Cat# 217604 |
| mProm-II™ Reverse Transcription System | Promega | Cat# A3800 |
| Nextera XT DNA Library preparation kit | Illumina | Cat# FC-131-1024 |
| P3 primary cell solution 4D-Nucleofector™ X Kit S | Lonza | Cat# V4XP-3032 |
| Platelet VASP/P2Y12 | Biocytex | REF #7014 |

| | | |
|---|---|---|
| PowerUp$^{TM}$ SYBR$^{TM}$ Green Master Mix for qPCR | Applied Biosystem | Cat# A25742 |
| qScript cDNA SuperMix | Quanta bio | Cat# 84034 |
| RNA 6000 Nano Kit | Agilent | Cat# 5067-1511 RUO |
| SMARTer Ultra Low Input RNA kit | Clontech | Cat# 634940 |
| TaqMan Advanced miRNA cDNA Synthesis Kit | Applied Biosystem | Cat# A28007 |
| TaqMan Fast Advanced Master Mix | Applied Biosystem | Cat# 4444557 |
| TaqMan$^{TM}$ MicroRNA Reverse Transcription Kit | Applied Biosystem | Cat# 4366596 |
| **Plasmids** | | |
| *Tol2-lyzC-RFP-miR-223* | Addgene | Cat# 97148 |
| *Tol2-itga2b:dsRED-pri-miR-223* | Fontana Lab - prepared by: Veronika Zapilko | |
| *pKE4-I-SceI-fabp10a:Xpt-β-cat, cryaa:Venus* | Addgene | Cat# 105127 |
| *Tol2-itga2b:mCherry* | Fontana Lab - prepared by: Paulina Ciepla | |
| *Tol2-BFP* | Gift from Julien Bertrand | |
| **Restriction enzymes** | | |
| XmaI | NEB | Cat# R0180 |
| ZraI | NEB | Cat# R0659 |
| DpnI | NEB | Cat# R0176 |
| MluI-HF | NEB | Cat# R3198 |
| NcoI-HF | NEB | Cat# R3193 |
| KpnI-HF | NEB | Cat# R3142 |
| AgeI-HF | NEB | Cat# R3552 |
| **Other reagents** | | |
| BD Vacutainer EDTA Tubes | Becton, Dickinson and Company | Unknown - study performed between June 2006 and December 2008 |

***Supplemental Table S 4*** *List of Primers, siRNAs and miRNA probes*

| **Reagent** | **Source** | | |
|---|---|---|---|
| **qPCR primers** | **Direction** | **Sequence** | |
| Hs-HB2M | forward | 5'-TGCTCGCGCTACTCTCTCTTT-3' | |
| Hs-HB2M | reverse | 5'-TCTGCTGGATGACGTGAGTAAAC-3' | |
| Hs-MASTL | forward | 5'-GCTTTCTCAAGGACTCGTATGCC-3' | |
| Hs-MASTL | reverse | 5'-CATTCCTGGGTTGGAGGCAACT-3' | |
| **qPCR probes** | **Source** | **Identifier** | **Assay Number** |
| hsa-miR-106a-5p, | Applied Biosystem | Cat# A25575 | 478225_mir |
| hsa-miR-106b-5p, | Applied Biosystem | Cat# A25575 | 478412_mir |
| hsa-miR-142-5p, | Applied Biosystem | Cat# A25575 | 477911_mir |
| hsa-miR-16-5p | Applied Biosystem | Cat# A25575 | 477860_mir |
| hsa-miR-17-5p, | Applied Biosystem | Cat# A25575 | 478447_mir |
| hsa-miR-484 | Applied Biosystem | Cat# A25575 | 478308_mir |
| hsa-miR-93-5p, | Applied Biosystem | Cat# A25575 | 478210_mir |
| **RT and qPCR probes** | | | |
| U6 snRNA | Applied Biosystem | Cat# 4427975 | 001973 |
| dre-miR-150 | Applied Biosystem | Cat# 4440886 | 006027_mat |
| hsa-miR-126 | Applied Biosystem | Cat# 4427975 | 000450 |
| hsa-miR-223 | Applied Biosystem | Cat# 4427975 | 000526 |
| **miRNAs** | | | |
| cel-miR-39-3p mirVana miRNA mimic | Ambion | Cat# 4464066 | MC10956 |
| **siRNAs** | **Source** | **Identifier** | **Sequence** |
| Hs_MASTL_7 | Qiagen | Cat# SI02653182 | 5'-CAGGACAAGTGTTATCGCTTA-3' |
| AllStars Neg. Control siRNA | Qiagen | Cat# 1027281 | undisclosed |
